# How much fear is in anxiety?

**DOI:** 10.1101/385823

**Authors:** Andreas J. Genewsky, Nina Albrecht, Simona A. Bura, Paul M. Kaplick, Daniel E. Heinz, Markus Nußbaumer, Mareen Engel, Barbara Grünecker, Sebastian F. Kaltwasser, Caitlin J. Riebe, Benedikt T. Bedenk, Michael Czisch, Carsten T. Wotjak

## Abstract

The selective breeding for extreme behavior on the elevated plus-maze (EPM) resulted in two mouse lines namely high-anxiety behaving (HAB) and low-anxiety behaving (LAB) mice. Using novel behavioral tests we demonstrate that HAB animals additionally exhibit maladaptive escape behavior and defensive vocalizations, whereas LAB mice show profound deficits in escaping from approaching threats which partially results from sensory deficits. We could relate these behavioral distortions to tonic changes in brain activity within the periaqueductal gray (PAG) in HAB mice and the superior colliculus (SC) in LAB mice, using in vivo manganese-enhanced MRI (MEMRI) followed by pharmacological or chemogenetic interventions. Therefore, midbrain-tectal structures govern the expression of both anxiety-like behavior and defensive responses. Our results challenge the uncritical use of the anthropomorphic terms *anxiety* or *anxiety-like* for the description of mouse behavior, as they imply higher cognitive processes, which are not necessarily in place.

## Introduction

The anthropomorphic terms *anxiety* or *anxiety-like* are widely used for the description of affective states in laboratory animals. The definition for anxiety (*American Psychiatric Association, 2013*) includes worries about distant or potential threats while the occurrence of exaggerated anxiety in combination with constant ruminations about illusionary threats indicates an anxiety disorder. *Fear* on the other hand describes the affective state (*‘being afraid’*) which is elicited with respect to an explicit, threatening stimulus.

The behavioral repertoire of fear - i.e. the sum of defensive responses - results from a recruitment of the *defensive survival circuits* (*LeDoux, 2014*). Its functions are either increasing the distance between the subject and the threat (flight), rendering the subject invisible to the threat (freezing) or ultimately enabeling the subject to fight. This includes the autonomic and neuroendocrine processes which prepare the creature for a successful flight e.g. reflected by increased heart and respiratory rate and release of stress hormones via increased hypothalamus-pituitary-adrenal-medulla (HPA) axis activity. As previously suggested, this condition is described best as the *defensive organismic state* (*LeDoux, 2014*). Therefore, it is just to say that the subjective feeling of being anxious or afraid are cognitive processes, while the behavioral expression of anxiety, fear and panic are physical or bodily processes which are typically orchestrated by subcortical and mesencephalic structures (*LeDoux and Pine, 2016*). In laboratory animals, like mice and rats, we lack the access to these subjective inner cognitive states, but have to solely rely on the interpretation of physiological and behavioral data.

A variety of behavioral testing paradigms therefore aims to assess states of *anxiety, fear* or *panic* based on the type and quality of evoked defensive behaviors in response to specific stimuli or contexts (*for review see Cryan and Holmes, 2005; Calhoon and Tye, 2015*). Hereby, more subtle be haviors like avoiding exposed and brightly illuminated areas on an elevated plus maze (EPM)(*Pellow et al., 1985*) are interpreted as *anxiety*. In contrast, the sudden jumping (a flight reaction completely different from startle response) followed by pronounced immobility (freezing) upon the onset of a previously negatively conditioned tone (auditory/Pavlovian fear conditioning; *for review see Maren, 2001*) is commonly associated with *fear*. These tests suggest a sharp distinction between the behavioral measures of *anxiety* and *fear*. For instance, auditory fear conditioning experiments paved the way for an in depth understanding of the amygdalar circuits underlying the expression a single characteristic defensive response (i.e., freezing) (*for review see Tovote et al., 2015*). In more complex and ethological relevant testing situations, however, one can observe a gradual transition from risk assessment to avoidance and flight or tonic immobility and ultimately fight/panic-like jumping as a function of the threat’s imminence (i.e. defensive distance) and the ability to escape (*Ratner, 1967,1975; Blanchard et al., 1986; Blanchard and Blanchard, 1990; Blanchard et al., 1990, 1997,2003*). This relationship was initially conceptualized as the *predatory imminence continuum (Fanselow and Lester, 1988*) and later has been integrated into the *two-dimensional defense system (McNaughton and Corr, 2004*). The two-dimensional defense system is of particular significance as it comprehensively describes the interplay of *defensive avoidance* and *defensive approach* with respect to the *defensive distance* (perceived distance to threat). In addition, it highlights the functional hierarchy of dominant brain structures in the orchestration of the behavioral expression of anxiety, fear and panic. In this context, McNaughton & Corr reappraise the function of the periaqueductal gray (PAG) *‘in the lowest levels of control of anxiety’ (McNaughton and Corr, 2004) (see also (McNaughton and Corr, 2018)*).

In this line of thinking we were interested to which extent the behavioral phenotype of a mouse model for extremes in *trait anxiety* (**1**) is accompanied by altered levels of defensive responses, and in addition (**2**) can be explained by changed neuronal activity in midbrain structures. As a model organism we chose two mouse lines which were previously established from CD1 mice as the result of a selective breeding approach based on the behavior on the EPM - a classical anxiety test. Thereby hyperanxious high-anxiety behaving (HAB) and hypoanxious low-anxiety behaving (LAB) mice were generated (*Krömer et al., 2005*) which are compared to normal-anxiety behaving (NAB) mice. Besides the already mentioned *anxiety-like* phenotype on the EPM (*Krömer et al., 2005; Bunck et al., 2009; Erhardt et al., 2011; Avrabos et al., 2013; Yen et al., 2013; Füchsl et al., 2014*), these lines show also marked differences in other behavioral and physiological measures (*see* Table 1). In HAB mice, most of the behavioral measures are biased towards immobility or lack of exploratory drive. This bears the risk of false interpretations, since altered locomotor activity and/or motivation might explain the extreme phenotypes as well. In the present study we comprehensively re-characterize HAB, NAB and LAB (HNL) mice for their extreme behavioral phenotypes on the EPM. We provide evidence that in HAB animals only ethobehavioral EPM measures and the levels of autonomic arousal are sensitive to anxiolytic treatment. In addition, we demonstrate for the first time that adult HAB animals show a disposition for sonic/audible vocalizations which is decreased by the anxiolytic diazepam. Further, we show that the extremes in high or low *anxiety-like* behavior of HAB and LAB animals are accompanied by paralleled alterations active in defensive responses using two novel, multi-sensory tasks (Robocat and IndyMaze) which assay repeated, innate escape behavior towards an approaching threatening stimulus. Hereby, we demonstrate that HAB animals present maladaptively altered levels of defensive responses, while LAB animals exhibit a strongly deficient reaction towards the threatening stimulus. Using several complementary strategies to probe the visual capabilities of HNL animals (optomotor response, electroretinography, etc.), we show that LAB animals suffer from complete retinal blindness. In order to assess tonic/basal in-*vivo* whole-brain neuronal activity alterations in HAB and LAB animals, we employ manganese-enhanced magnetic resonance imaging (MEMRI) (*Grünecker et al., 2010; Bedenk et al., 2018*). Thereby, we provide evidence that HAB mice exhibit an increased neuronal activity within the PAG, while LAB mice show a decreased activity in the deep layers of the superior colliculus (SC). Finally, using a designer receptor exclusively activated by designer drugs (DREADD) approach in LAB mice or by applying localized injections of muscimol in HAB mice we are able to partially revert the extreme phenotypes in *anxiety-like* behavior in LAB and HAB animals.

**Table 1.**
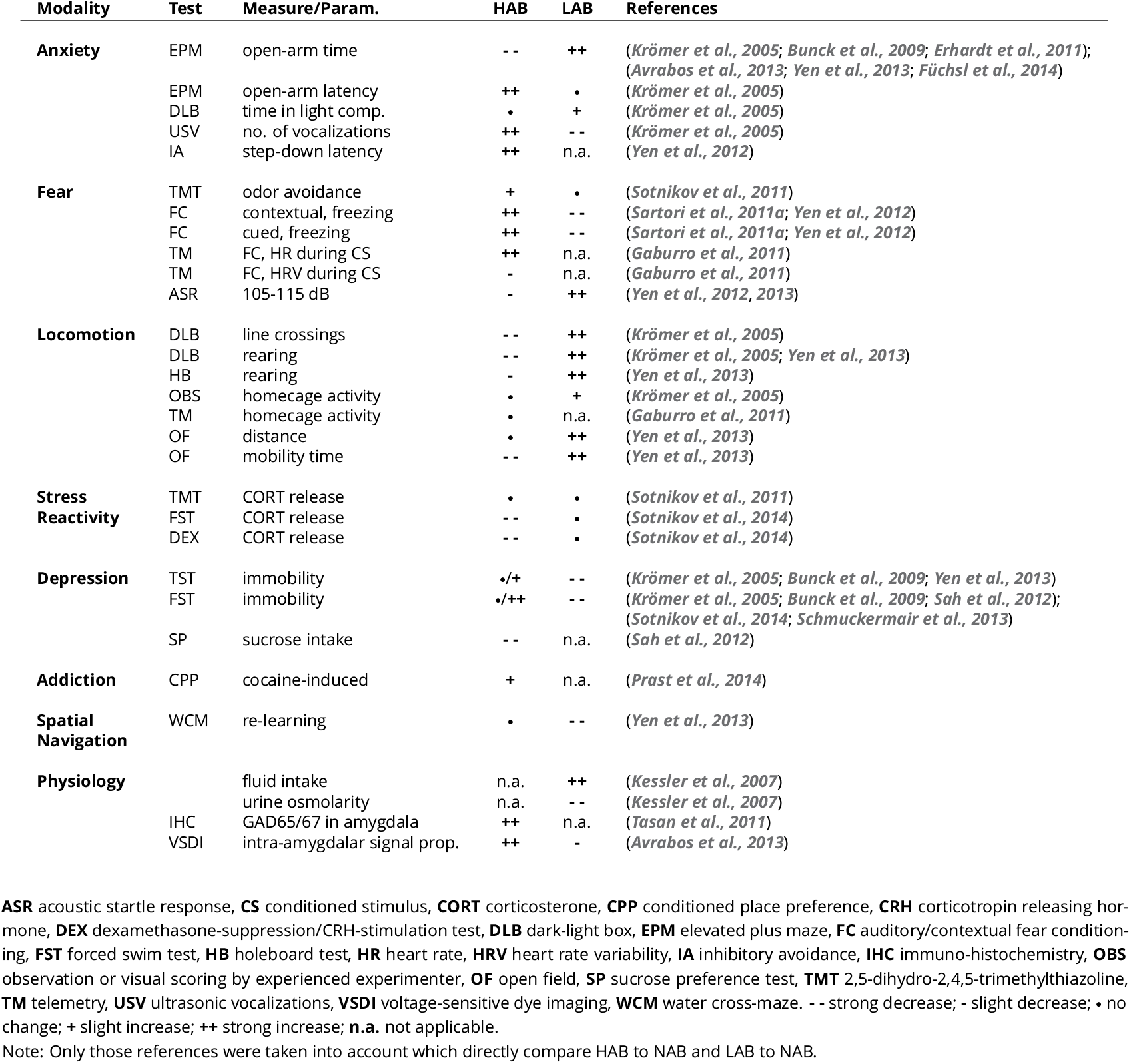
Physiological & Behavioral Phenotypes of HAB and LAB mice

| Modality | Test | Measure/Param. | HAB | LAB | References |
| --- | --- | --- | --- | --- | --- |
| **Anxiety** | EPM | open-arm time | -- | ++ | (Krömer et al., 2005; Bunck et al., 2009; Erhardt et al., 2011); (Avrabet et al., 2013; Yen et al., 2013; Füchsl et al., 2014) |
|  | EPM | open-arm latency | ++ | • | (Krömer et al., 2005) |
|  | DLB | time in light comp. | • | + | (Krömer et al., 2005) |
|  | USV | no. of vocalizations | ++ | -- | (Krömer et al., 2005) |
|  | IA | step-down latency | ++ | n.a. | (Yen et al., 2012) |
| **Fear** | TMT | odor avoidance | + | • | (Sotnikov et al., 2011) |
|  | FC | contextual, freezing | ++ | -- | (Sartori et al., 2011a; Yen et al., 2012) |
|  | FC | cued, freezing | ++ | -- | (Sartori et al., 2011a; Yen et al., 2012) |
|  | TM | FC, HR during CS | ++ | n.a. | (Gaburro et al., 2011) |
|  | TM | FC, HRV during CS | - | n.a. | (Gaburro et al., 2011) |
|  | ASR | 105-115 dB | - | ++ | (Yen et al., 2012, 2013) |
| **Locomotion** | DLB | line crossings | -- | ++ | (Krömer et al., 2005) |
|  | DLB | rearing | -- | ++ | (Krömer et al., 2005; Yen et al., 2013) |
|  | HB | rearing | - | ++ | (Yen et al., 2013) |
|  | OBS | homecage activity | • | + | (Krömer et al., 2005) |
|  | TM | homecage activity | • | n.a. | (Gaburro et al., 2011) |
|  | OF | distance | • | ++ | (Yen et al., 2013) |
|  | OF | mobility time | -- | ++ | (Yen et al., 2013) |
| **Stress Reactivity** | TMT | CORT release | • | • | (Sotnikov et al., 2011) |
|  | FST | CORT release | -- | • | (Sotnikov et al., 2014) |
|  | DEX | CORT release | -- | • | (Sotnikov et al., 2014) |
| **Depression** | TST | immobility | •/+ | -- | (Krömer et al., 2005; Bunck et al., 2009; Yen et al., 2013) |
|  | FST | immobility | •/++ | -- | (Krömer et al., 2005; Bunck et al., 2009; Sah et al., 2012); (Sotnikov et al., 2014; Schmuckermair et al., 2013) |
|  | SP | sucrose intake | -- | n.a. | (Sah et al., 2012) |
| **Addiction** | CPP | cocaine-induced | + | n.a. | (Prast et al., 2014) |
| **Spatial Navigation** | WCM | re-learning | • | -- | (Yen et al., 2013) |
| **Physiology** |  | fluid intake | n.a. | ++ | (Kessler et al., 2007) |
|  |  | urine osmolarity | n.a. | -- | (Kessler et al., 2007) |
|  | IHC | GAD65/67 in amygdala | ++ | n.a. | (Tasan et al., 2011) |
|  | VSDI | intra-amygdalar signal prop. | ++ | - | (Avrabet et al., 2013) |

**ASR** acoustic startle response, **CS** conditioned stimulus, **CORT** corticosterone, **CPP** conditioned place preference, **CRH** corticotropin releasing hormone, **DEX** dexamethasone-suppression/CRH-stimulation test, **DLB** dark-light box, **EPM** elevated plus maze, **FC** auditory/contextual fear conditioning, **FST** forced swim test, **HB** holeboard test, **HR** heart rate, **HRV** heart rate variability, **IA** inhibitory avoidance, **IHC** immuno-histochemistry, **OBS** observation or visual scoring by experienced experimenter, **OF** open field, **SP** sucrose preference test, **TMT** 2,5-dihydro-2,4,5-trimethylthiazoline, **TM** telemetry, **USV** ultrasonic vocalizations, **VSDI** voltage-sensitive dye imaging, **WCM** water cross-maze. -- strong decrease; - slight decrease; • no change; + slight increase; ++ strong increase; n.a. not applicable.
Note: Only those references were taken into account which directly compare HAB to NAB and LAB to NAB.

## Results

### Behavioral Assessment of HAB, NAB, LAB mice on the Elevated Plus Maze

The elevated-plus maze (EPM) is considered to be a robust assay for the detection of altered *anxiety-like* behavior in mice. However, the standard test duration rarely exceeds 5-10 minutes (*Komada et al., 2008*), whereby strong inter-individual differences in avoidance behavior and especially their pharmacological modulation, are masked due to stringent cut-off criteria. In order to overcome this issue, we have extended the testing duration to 30 minutes and re-evaluated the behavior of HAB (*N*=11), NAB (*N*=7) and LAB (N=7) mice on the EPM, while focusing on the initial 5 minutes for all parameters, except for latency (0-30 min) and stretch-attend postures (0-15 min), to provide measures which are largely comparable to previous studies (*see* Fig 1A). Analysis of data obtained during the entire observation period revealed essentially the same findings (not shown).

Using this approach, significant group differences (*F*_2,22_=15.07, *p*<0.0001) in the latencies to explore the open arms were revealed (Fig 1A). More than 45% of all HAB animals did not enter the open arm, even within the extended testing duration of 30 minutes compared to 0% in NAB mice (*χ*^2^=4.41, *p*=0.0358). On the contrary all LAB animals explored the open arm with latencies < 6 minutes. These distinct behavioral traits were also reflected by the percentage of time the animals spent on the open arm: LAB animals 53.6±11.3% compared to 2.4±0.8% NAB (*F*_2,22_ =26.25, *p*<0.0001). Additionally, LAB animals showed an overall increase in locomotor activity (1400.0±171.7 cm vs. 723.0±60.8 cm, *F*_2,22_=22.49, *p*<0.0001). On the contrary, HAB animals spent more than 85% of the time in the closed arm (*F*_2,22_=28.98, *p*<0.0001), as they also avoided staying in the central zone (13.0±2.3 % vs. 33.8±4.0 %, *F*_2,22_=12.96, *p*=0.002). These observations are consistent with previous reports of HAB, NAB and LAB behavior on the EPM (*Krömer et al., 2005; Bunck et al., 2009; Erhardt et al., 2011; Avrabos et al., 2013; Yen et al., 2013; Füchsl et al., 2014*). The rather low open-arm time shown by NAB mice may relate to the specific test conditions (we placed the EPM in middle of a large, dimly lit room without additional surrounding enclosures). To complement the traditional EPM parameters, the display of stretched-attend postures (SAP) (*Grant and Mackintosh, 1963*), a form of active risk assessment behavior, was analyzed as an ethobehavioral measure (Fig 1B). It was previously shown that the number of SAPs decreases upon anxiolytic treatment (*Kaesermann, 1986*) and increases with the anxiogenic 5-HT_2C/1B_ receptor antagonist mCPP (*Grewal et al., 1997*). Moreover, the display of SAPs depend on the presence of an imminent threat or a potential threatening situation and demonstrate the general motivation of the animals to explore a potentially threatening environment (*Pinel et al., 1989*). LAB (*N*=6, one animal was excluded as no SAPs were displayed) animals showed a significantly lower number of SAPs (Fig 1B; *F*_2,19_=29.84, *p*<0.0001; LAB 20.7±5.9 vs. NAB 63.0±4.3), whereas HAB animals were indistinguishable from NAB (Fig 1B; *N*=5, two animal were excluded as no SAPs were displayed). Looking at the overall duration of displayed SAPs, HAB animals showed increased measures (HAB 222.4±16.8 s vs. NAB 158.0±15.2 s), whereas LAB animals spent on average only 33.7±12.3 seconds displaying SAPs (*F*_2,19_=32.74, *p*<0.0001). If analyzed in 5 min bins, NAB animals could adapt to the EPM and the duration of displayed SAPs decayed. On the contrary, HAB animals showed an elevated non-decaying response after 15 minutes (group×time interaction: *F*_2,28_=3.587, *p*=0.0410; 2-way rmANOVA) and higher autonomic arousal, which was reflected by significantly increased defecation (*Hall, 1934*) during the EPM task (Fig 1C; 11.6±1.2 vs. 7.7±0.7, *F*_2,21_ = 4.779, *p*<0.0195).

Before the animals were placed on the EPM, every subject was tested for the disposition to emit sonic vocalizations by lifting them 3 times from grid cage top (*Whitney, 1970*). Animals which vocalized at least once, were counted as ‘vocalizers’. Whereas none of the NAB (*N*=13) or LAB (*N*=15) animals emitted even a single call, 47% of HAB (*N*=15) animals vocalized at least once (Fig 1D; *χ*^2^=15.61, *p*=0.0004).

In order to investigate to which extent the phenotype of HAB mice can be modulated with traditional anxiolytics, we injected diazepam (DZP, 1 mg/kg i.p.) or vehicle (saline) to separate groups of experimentally naive HAB mice (*N*=13, each). None of the classical EPM parameters were sensitive to DZP treatment, except for an increase in locomotor activity (Fig 1E; t_24_=2.174, *p*=0.0398). Neither the total number nor the total duration of SAPs were significantly altered by DZP (Fig 1F). However, analysis in 5-min bins revealed that DZP turned the non-decaying display of SAPs shown by vehicle-treated HAB into a decaying trajectory (treatment×time: interaction: *F*_2,28_=3.587, *p*=0.0410; 2-way rmANOVA) which resembles the situation in NAB mice. In addition, DZP treatment decreased defecation (Fig 1G; 1.5±0.5 vs. 5.8±1.2, t_24_=3.344, *p*=0.0027) and the disposition to vocalize during a 5 minute tail-suspension test(TST) (Fig 1H; 3 out of 13 vs. 9 out of 13, two-sided Fisher’s exact test *p*=0.0472). The higher absolute incidence of vocalizers, compared to the data shown in Fig 1D, is most likely due to prior injection stress. The lower absolute defecation scores, in turn, might be partially ascribed to defecation during the injection procedure. Taken together, HAB, NAB and LAB animals show a robust behavioral phenotype on the EPM. Further, under our experimental conditions, the traditional EPM measures are not sensitive to diazepam-treatment, but more ethologically relevant measures like autonomic arousal, vocalization and active risk assessment.

### Two Novel Ethologically Inspired Testing Situations Reveals Extremes in Innate Defensive Responses in HAB and LAB Mice

The behavioral measures obtained on the EPM are indicative of an approach-avoidance situation which became manifest differently in HAB and LAB mice. The term *anxiety* test for the EPM infers an inner conflict which misleadingly points towards higher cognitive processes, mediated for example by the prefrontal areas. Looking at avoidance behavior separately, it becomes obvious that there is a strong subcortical component which is in a continuum to flight and *panic-like* reactions, involving most likely the amygdala, ventromedial hypothalamus, periaqueductal gray and the superior colliculus (*McNaughton and Corr, 2018*). Therefore we were interested if the altered EPM phenotype of HAB and LAB is accompanied by changes in defensive responses as it has been suggested previously to be the case with conditioned fear (*Sartori et al., 2011b; Yen et al., 2012*). In order to circumvent learning mediated effects, we focused on innate defensive responses upon acute confrontation with a (potential) threat.

Paradigms which asses general innate *fear* levels should incorporate multi-sensory stimuli and allow for repeated testing and temporally confined exposure. In lack of appropriate testing situations, we have developed two novel paradigms: the Robocat (*see*), which is based on a previously published design by *Choi and Kim (2010*), and the IndyMaze, which is inspired by a popular movie (*Spielberg and Marshall, 1981*) (For a detailed description of both tests *see* section *Methods and Materials*). The different behavioral readouts obtained in the Robocat task are depicted in Fig 2A. The mouse could either activate the Robocat and subsequently display a flight response, activate the Robocat but simply bypassing it or activate the Robocat and collide with it. The innate defensive responses of HAB (*N*=7), NAB (*N*=6) and LAB (*N*=9) mice were assessed using the Robocat task. Fig 2B depicts the percentage of animals which displayed the respective behaviors at least once during a 10 minute exposure to the Robocat. During this trial the animals activated the Robocat several times (HAB 2.4±0.4, NAB 3.5±0.6, LAB 10.8±2.1). HAB animals were not able to adapt to the Robocat’s activation and showed a flight response at all encounters (Fisher’s exact *p*=0.021), they never bypassed (Fisher’s exact *p*=0.0047) nor collided with it. On the contrary NAB animals, displayed a well-balanced behavioral profile: the minority of all animals fled the Robocat (33%) or got hit by the it (17%), while 83% of all NAB mice tolerated and bypassed the threatening stimulus at least once. This is contrasted by the behavior of LAB mice: no single animal fled upon the Robocat’s movement, but all bypassed it. Most strikingly however, the vast majority of LAB mice 89% even collided with it at least once (Fisher’s exact *p*=0.011).

**Figure 1.**
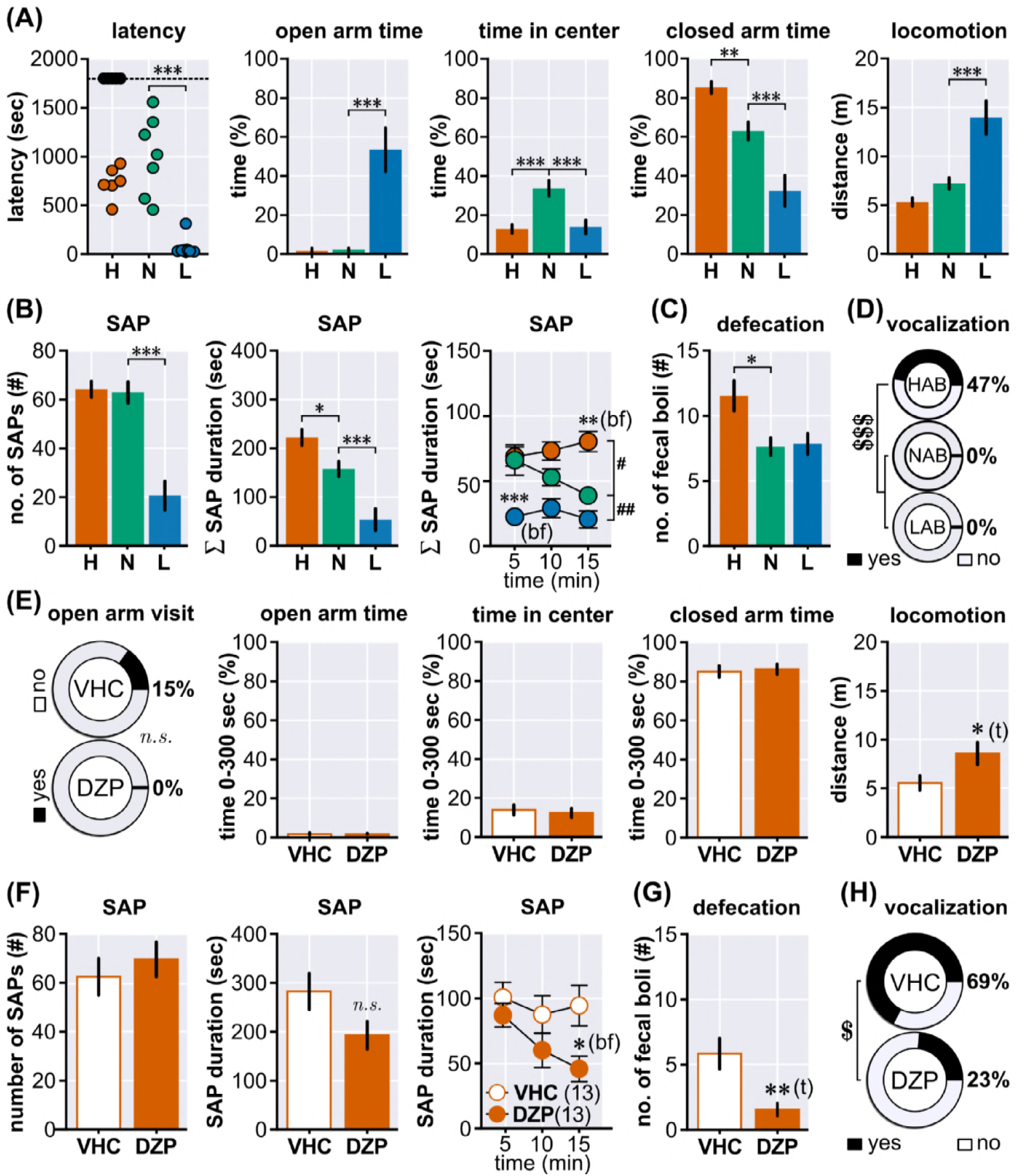
Behavioral Assessment of HAB, NAB, LAB Mice on the EPM. (**A**) Using the standard EPM but with an extended cut-off time of 30 min the following behavioral parameters were assessed for HAB (*N*=11), NAB (*N*=7) and LAB (*N*=7) animals: the latency to enter the open arm, open-arm time (first 5 minutes; 0-5 min), central zone time (0-5 min), closed arm time (0-5 min) and the distance the animals have traveled (0-5 min). (**B**) In addition to the classical EPM parameters we have also investigated the display of stretched-attend postures (SAP) which serves as a measure of active risk assessment: the total number of SAPs during the first 15 min of the task, the total duration of SAPs during the first 15 min, and the duration of SAPs in 5 minute bins. (**C**) Defecation during EPM exposure (number of fecal boli) as an indirect measure of autonomic arousal. (**D**) Disposition to emit sonic/audible vocalizations. (**E**) A new cohort of experimentally naïve HAB mice was treated with diazepam (1 mg/kg, i.p.;*N*=13) or vehicle (saline, *N*=13) before exposure to the EPM. (**F**) Stretched-attend posture display of HAB animals during EPM with diazepam/vehicle treatment. (**G**) Defecation of HAB mice. (**H**) In order to assess the disposition to vocalize in standardized manner, the diazepam/vehicle treated HAB animals were subjected to a 5 min tail-suspension test, while audio signal were recorded and scored offline. Asterisks indicate significance values obtained by *t*-tests (t) or 1-way ANOVA followed by Newman-Keuls Multiple Comparison/Bonferroni post-hoc (bf) tests, \**p*<0.05, ** *p*<0.01, *** *p*<0.001; dollar signs indicate significance values obtained by *χ*^2^ tests, $ *p*<0.05, $$$ *p*<0.001; hashes indicate group effects obtained by 2-way ANOVA, ## *p*<0.01. Values are given as mean±SEM.

The Robocat task revealed differential defensive responses between HAB and NAB, whereby the inability of HAB mice to bypass the Robocat can be interpreted as maladaptive behavior. At the same time the high degree of controllability (allowing a bypass or withdrawal from the arena to avoid activation/confrontation) does not allow to ask whether NAB and LAB show different levels of defensive responses: the inability to express defensive responses, and a high degree of adaptation would result both in a decreased level of observable defensive reactions. In order to avoid this confounding variable we have developed the IndyMaze. In this test an animal is confronted with a rolling (25 cm/s) styrofoam ball (100 g) in a tilted (<1 °) and narrow tunnel. Therefore, every trial involves a direct encounter with the threatening stimulus. The operational procedure is depicted in Fig 2C. First, the animals are free to enter the arena, which gives the latency to first exit, a measure comparable to other emergence tasks (Fig 2D). This measure corresponds to the exit latency on the EPM. HAB animals showed high latencies to exit the home compartment (HAB 977.4±79.2 s vs. NAB 392.1 ±66.1 s) whereas LAB animals were not different from NAB (*F*_2,50_=12.64, *p*<0.0001). A significant amount of HAB animals never left the home compartment (Fig 2E) within 30 minutes (*χ*^2^_2,*N*=62_=6.671, *p*=0.0356). Once the animals have left the home compartment, they explored the entire arena (Fig 2F) with equally low latency (HAB 68.8±10.1 s; NAB 213.1 ±72.8 s; LAB 100.4±18.2 s). This demonstrates comparable levels of exploratory drive in all three lines and precludes that the increased latency to the 1^st^ exit simply results from a lack of motivation or impaired locomotor behavior. Looking at the defensive responses (Fig 2G), which included preemptive flight responses or a retrieval after the ball has hit the animals, it is evident that both HAB and NAB are able to respond appropriately towards approaching threatening stimuli, whereas 60% of LAB animals exhibited significant deficits and failed to generate at least one defensive reaction (*χ*^2^_1*N*=27_=13.11, *p*=0.0014). In order to test whether the behavioral readouts obtained using the IndyMaze can be modulated with anxiolytics, another cohort of HAB animals was treated with diazepam (DZP, 1 mg/kg, *N*=13) or vehicle (VHC, saline, *N*=12) and were subjected to the IndyMaze task. The DZP treatment could significantly decrease the latency to 1^st^ exit (Fig 2H; VHC 1011.0±153.4 s vs. DZP 595.5±133.5 s; Mann-Whitney, two-tailed, *U*_n1=208, n2=117_=39.00, *p*<0.0363), indicative of an anxiolytic effect, while leaving latency for end-exploration unaffected (*see*). However, DZP treatment was ineffective in modulating defensive responses (defensive responsivity: VHC 100%, DZP 100%). NAB and HAB, but not LAB, mice showed short-term avoidance of additional encounters with the ball, as indicated by the increase in latency until re-entering the middle part of the arena. One week later, both HAB and NAB mice showed a highly significant decrease in latency to 1st entry compared to the first exposure. Nevertheless, only NAB mice showed long-term avoidance of the middle segment of the arena, which is indicative of maladaptive consequences of heightened fear/ anxiety for the development of avoidance behavior (data not shown).

In summary, both tasks, the Robocat and IndyMaze, have proven to be valid tools to assay innate defensive responses in mice. In addition, the IndyMaze task permits also the parallel assessment of inhibitory avoidance behavior. Using both tasks, we could demonstrate that HAB mice show maladaptive levels of defensive responses. LAB animals, in contrast, exhibited strong deficits to escape imminent threats.

### Complete Retinal Blindness in LAB Mice

The remarkable ignorance of LAB mice to approaching objects forced us to look for differences in visual perception. A standard test for visual acuity in mice is the assessment of the optomotor response (OMR) (*Thaung et al., 2002; Abdeljalil et al., 2005*). This test is based on the tracking behavior of mice in response to horizontally moving stripes. For this test, mice are placed on a fixed platform within a rotating cylinder lined with stripes of different width to probe visual acuity (Fig 3A *inset*). We modified this testing procedure in order to fit to all five mouse lines (B6, CD1, HAB, NAB and LAB) in a way that we have used only one, relatively large grating (0.5 cycles/degree) and in addition scored every head movement if it was concordant with the cylinders rotational direction.

**Figure 2.**
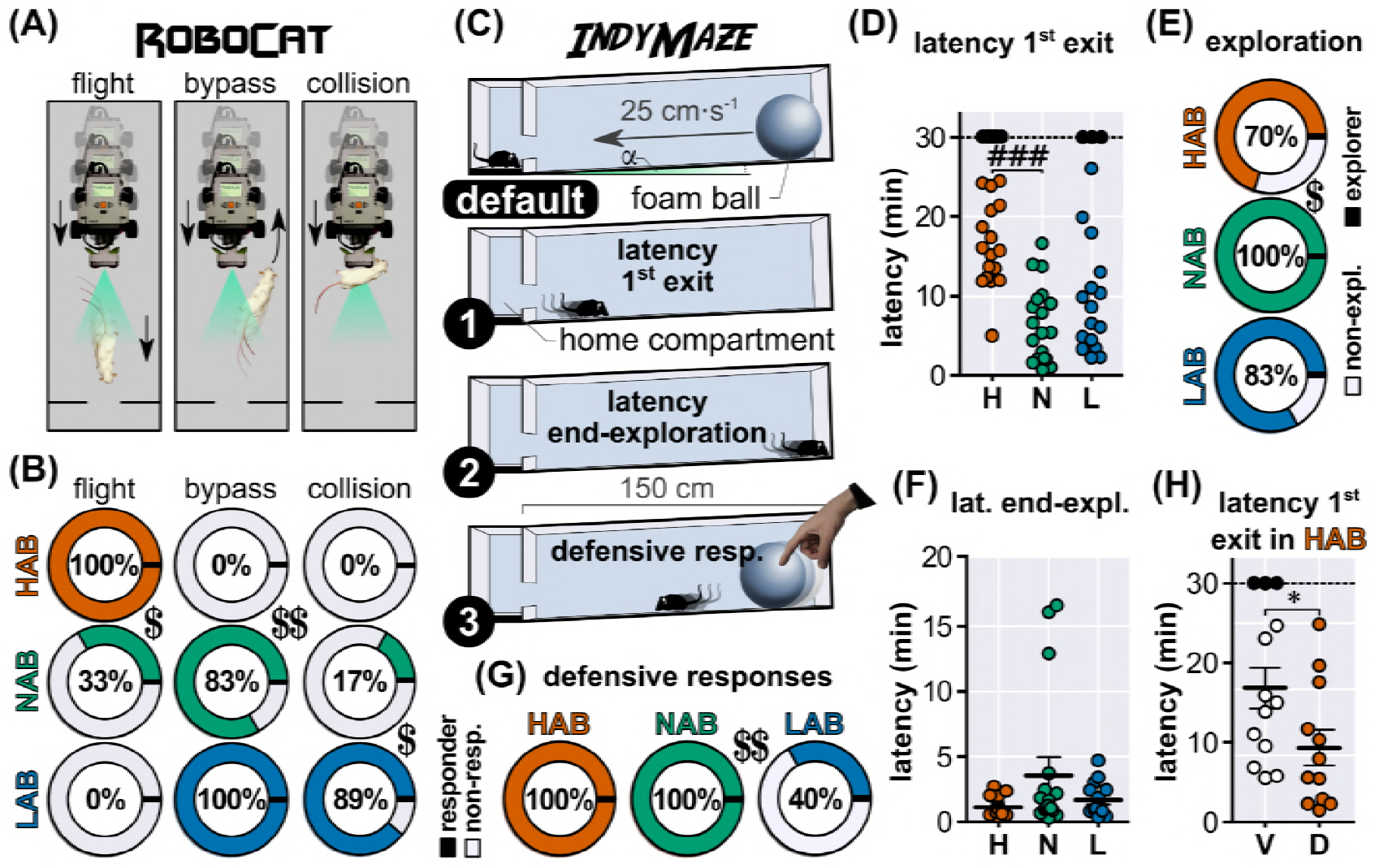
Two Novel Ethologically Inspired Testing Situations Reveals Extremes in Innate Defensive Responses in HAB and LAB Mice **A**) The three different behavioral measures obtained in the Robocat task, whose appearance have been scored: flight, bypass and collision. (**B**) Using the Robocat, we have investigated the fear responses of HAB (*N*=7, red), NAB (*N*=6, green) and LAB (*N*=9, blue) animals (analyzed using Fisher’s exact tests). Values are percentages of animals which showed the respective behavior at least once. (**C**) Schematic description of the IndyMaze (default) operational procedure as well as the different behavioral measures (latency to 1^st^ exit I, latency for end-exploration II and flight response III). Using the IndyMaze we have tested different cohorts of HAB (*N*=24), NAB (*N*=19) and LAB (*N*=20) animals. (**D**) Quantification of the latency to 1^st^; black-filled circles indicate animals which did not leave the start arm, HAB (*N*=7), NAB (*N*=0) and LAB (*N*=4), those animals were excluded from the 1-way ANOVA. (**E**) Quantification of the number of animals which explored the arena. (**F**) Quantification of the latency for end-exploration excluding animals which did not enter the arena at all, as shown in *D*. Note: If the animals left the start compartment, they all explored the arena to its end with comparable vigor. (**G**) Quantification of the occurrence of fear responses at least once during 3 encounters of the approaching styrofoam ball (this includes preemptive fear responses, as well as fear responses after the ball had hit the animal). (**H**) Another cohort of HAB animals was treated with diazepam (DZP, 1 mg/kg, i.p.) (*N*=13) or vehicle (VHC, saline, *N*=12) and subjected to the IndyMaze, and the latency to 1^st^ exit was quantified. Asterisks indicate significance values obtained by Mann-Whitney test, * = *p*<0.05;dollar signs indicate significance values obtained by *χ*^2^ or Fisher’s exact tests, $ *p*<0.05, $$ *p*<0.01, $$$ *p*<0.001;hashes indicate significance values obtained by 1-way ANOVA followed by Newman-Keuls Multiple Comparison test, ### *p*<0.001. Values are given as mean±SEM.

Therefore we have used this test to assess vision in general, rather than visual acuity. Using this approach, we could observe significant strain differences (Fig 3A; *F*_4,54_=93.13, *p*<0.0001). B6 mice outperformed all other strains by far (B6 16.9±0.9 OMR/min), whereas among the albino animals HAB animals showed the strongest responses (5.5±0.8 OMR/min). Both, CD1 (2.6±0.7 OMR/min) and NAB (2.4±0.5 OMR/min) animals responded similar, but LAB animals failed to show any clear optomotor responses (0.2±0.1 OMR/min).

As LAB mice also have been reported to exhibit certain phenomenological similarities to ADHD (Yen *et al., 2013*), we cannot exclude the possibility that these animals perceive but are unable to attend to the visual stimuli and thus fail to show an appropriate response. Therefore, the retinal function of all five mouse strains was investigated using flash electroretinography (fERG) measurements in the anesthetized animal. The fERG setup (depicted in Fig 3B *top*) consisted of a differential amplifier usually used for in-*vivo* extracellular neural recordings (*Siegle et al., 2017*), whereby the reference electrode was placed on the shaded eye. The other eye was stimulated with a custom built miniature eyecup, equipped with a white LED, in combination with a custom built LED driver. This setup allowed the reliable detection of electroretinographic signals and the dissection of the b-wave component of fERG (Fig 3B *bottom*). The fERGs acquired in scotopic (dark-adapted for >3h) as well under photopic conditions at three different light flash intensities (0.23, 128 and 1.69 log photoisomerizations×rod^-1^ ×s^-1^), showed strong deflections for B6, CD1, HAB and NAB (*N*=6, each) animals (Fig 3C). However, in LAB animals (*N*=6) there was no detectable electrophysiological response (scotopic: Group *F*_4,25_=14.38, *p*<0.0001, Fig 3D; photopic: Group *F*_4,2_5=8.77, *p*=0.0001, 2-way rmANOVA; Fig 3E). To further determine the cause for the absence of electroretinographic responses, a histological analysis of retinal sections of all strains (*N*=3, each, right eye) was conducted and an absence of the outer nuclear layer (ONL) and the subjacent inner/outer segments (IS/OS) was observed in LAB animals (Fig 3F). As the founder strain for LAB animals (CD1) is known to exhibit incidences of a recessive *rd1* retinal degeneration (*Serfilippi et al., 2004*), we employed a polymerase chain reaction (PCR) genotyping screening for all strains (*N*=4, each, tail biopsy) (*Chang et al., 2013*). The test (Fig 3G) revealed that LAB animals exhibit a homozygous mutation in the *Pde6b*^rd1+/+^ allele which is indicative of the retinal degeneration 1 mutation which leads to blindness shortly after birth. Therefore, it is to conclude that LAB animals (tested at an age of 3-6 month of age) suffer from complete retinal blindness, which is the reason for the inability to escape approaching threatening stimuli, like the Robocat (Fig 2B). But blindness does not explain why still only 40% of LAB animals showed a flight response even after hit by the ball in the IndyMaze task (Fig 2G).

**Figure 3.**
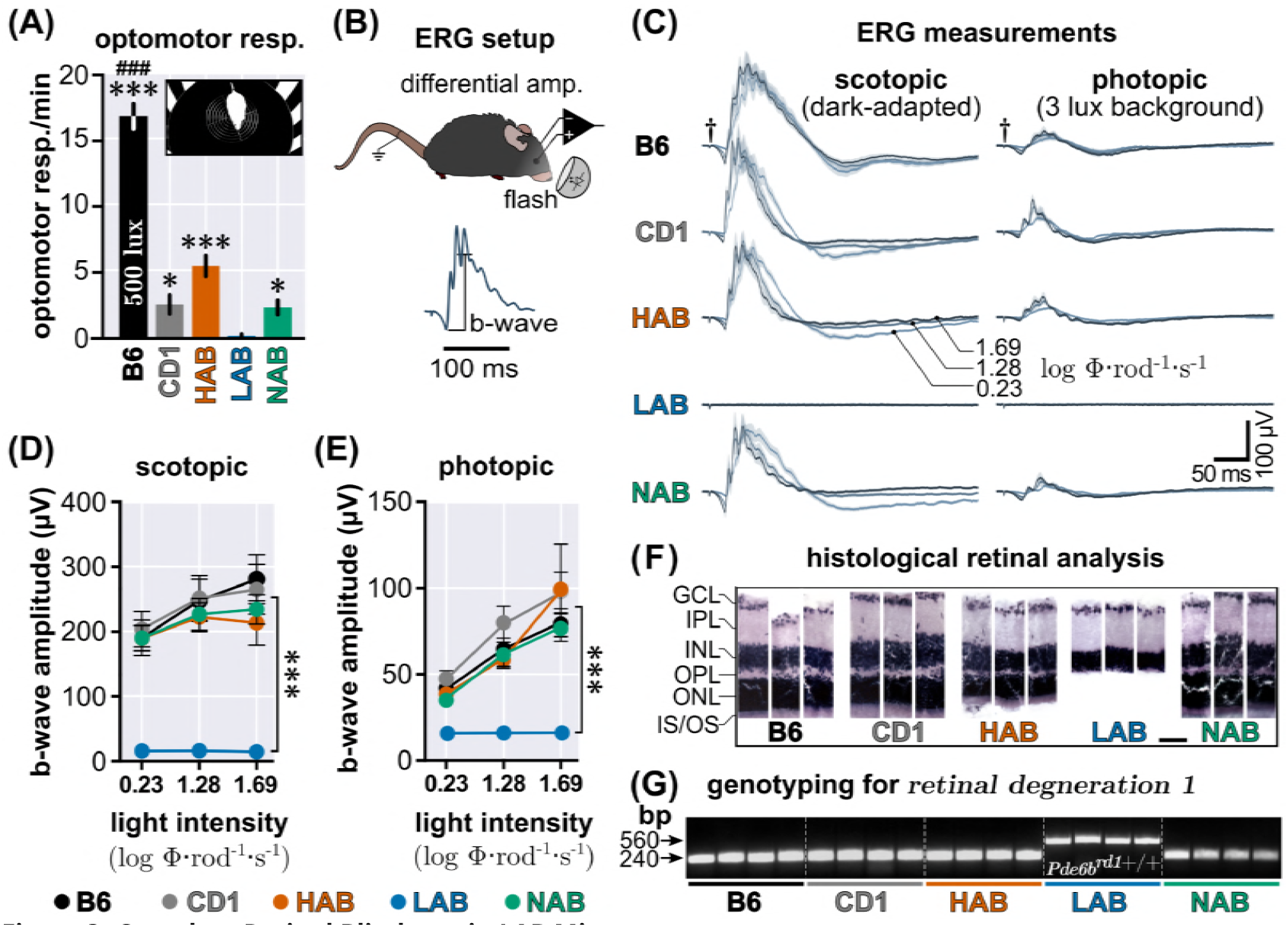
Complete Retinal Blindness in LAB Mice. (**A**) Optomotor responses (OMR) measured in B6, CD1, HAB, NAB, LAB (*N*=12, each) under 500 lux. Inset shows a HAB animal within the OMR setup. Significance values obtained by 1-way ANOVA followed by Newman Keuls Multiple Comparison are indicated by asterisks compared to LAB or by hashes for B6 compared to all other mouse lines. (**B**) Simplified overview of the setup for measuring electroretinography in the anesthetized mouse. The b-wave is typically associated with the activity of Müller and ON bipolar cells. (**C**) Electroretinograms of B6, CD1, HAB, NAB, LAB (*N*=6, each) measured at scotopic and photopic conditions a three different flash intensities. Quantification of (**D**) scotopic ERG and (**E**) photopic measurements. Asterisks indicate significant group effect obtained by 2-way ANOVA followed by Bonferroni post hoc test. (**F**) Histological analysis of 30 μm retinal sections of B6, CD1, HAB, NAB, LAB (*N*=3, each, right eye only) stained with haematoxylin and eosin. IS/OS inner/outer photoreceptor segments; ONL outer nuclear layer; OPL outer plexiform layer; INL inner nuclear layer; IPL inner plexiform layer; GCL ganglion cell layer. (**G**) Polymerase chain reaction (PCR) screening for *Pde6b*^rd1+/+^ allele, *retinal degeneration 1*; bp base pair. Significance values are indicated by asterisks and hashes (details for the statistical tests are given in the respective part of the figure legend): * *p*<0.05, ** *p*<0.01,*** *p*<0.001 vs. LAB; ### *p*<0.001 vs. CD1, HAB, LAB and NAB. Values are given as mean±SEM.

### Reversing the Low-anxiety Phenotype of LAB Mice

The severe deficit in avoiding approaching threats of LAB mice during the Robocat task is explained by their retinal degeneration. However, using the IndyMaze, where retrievals (after the ball had hit the animal) were also counted as fear responses, it became obvious that LAB mice also showed a decreased responsivity towards tactile stimuli. Therefore, it was necessary to determine whether this behavioral abnormality can be ascribed to differential activity in a certain brain area. In order to investigate the tonic neuronal activity changes in LAB mice compared to NAB, we employed manganese-enhanced magnetic resonance imaging (MEMRI) in HAB (*N*=31), NAB (*N*=26) and LAB (*N*=30) mice (FDR*p*<0.001, cluster extent >20), using a 3-level full-factorial design voxel-wise analysis 6 (for the complete MEMRI data set *see*). The results obtained by pairwise comparison of MEMRI data (i.e., HAB vs. NAB, LAB vs. NAB) suggested, among others, a decreased accumulation of manganese within the ventral parts of the deep and intermediate layers of the superior colliculus (lateral to the periaqueductal gray) in LAB mice (Fig 4A). This structure receives dense inputs from the primary and secondary somatomotor areas (Allen Brain Atlas, Connectivity, exps. #180719293, #180709942). In order to assess the functional relationship of this brain region in the generation of the LAB behavioral phenotype, the recently developed DREADD approach (*Armbruster et al., 2007*) was employed. The activating DREADD hM3Dq fused to the reporter protein mCherry was expressed under the control of the CaMKII*α* promoter using adeno-associated viral vectors (AAV5-CaMKII*α*-hM3Dq-mCherry, *N*=11) or the control virus (AAV5-CaMKIIα-mCherry, *N*=12) within the SC 6 (ML ±0.9 mm, AP −3.64 mm, DV −1.75 mm). An exemplary image of the virus expression is shown in Fig 4B. This approach resulted in the labeling of the entire SC (for detailed histological verification *see* A). After an incubation period of >5 weeks, all animals were subjected to the IndyMaze. On the testing day each animal was injected (i.p.) with 1 mg/kg CNO 45 minutes before the trial (as both, experimental and control animals, received the same amount of CNO, the previously discovered side-effects of converted clozapine (*Gomez et al., 2017*) cannot explain the behavioral changes). Experimental animals expressing hM3Dq showed a significantly (Mann-Whitney *U*_n1=94, n2=182_=16.00,*p*=0.0023) increased latency to leave the start compartment (1282.0±185.4 s vs. 265.5±113.6 s; Fig 4C). Moreover only 64% of hM3Dq animals left the start compartment (Fig 4D) within 30 minutes (Fisher’s exact, *p*=0.0373). Also the latency for end exploration was increased (216.0±62.7 s vs. 51.0±25.3 s; Mann-Whitney *U* test; *U*_n1=86, n2=104_=8.00, *p*=0.0046; Fig 4E). The fear responsivity (Fig 4F) was increased to 71% of mice transfected with hM3Dq, compared to 9% in mCherry controls (Fisher’s exact, *p*=0.0095). Next, we tested whether this pharmacogenetically augmented defensive response pattern, is also reflected by changed behavior on the EPM (one week after IndyMaze task, CNO injection 45 min prior to experiment; Fig 4G). Similar to the emergence component of the IndyMaze, hM3Dq animals treated with 1 mg/kg CNO showed an increased (85.6±12.9 s vs. 27.4±2.4 s) latency to access the open arm (*U*_n1=69.5, n2=183.5_ =3.500, *p*=0.0002). This was accompanied by a decreased percentage of time spent on the open arms (61.9±4.2 % vs. 4 76.0±3.3 %, *U*_n1=183, n2=93_=27.00, *p*=0.0178), an increased percentage of time spent in the closed 5 arms (33.6±4.3 % vs.18.3±3.0, *U*_n1=100.5, n2=17.5_= 22.50, *p*=0.0081) as well as decreased locomotor activity (7.1 ±0.7 m vs. 9.7±1.0 m, *U*_n1=179, n2=97_=31.00, *p*=0.0337). Time in center was unaffected (*see* B). The partially reverted behavioral phenotype of LAB mice on the EPM could be explained by an increased passivity due to nonspecific effects of the active DREADD. However, the increased number of active risk assessment behavior (Fig 4H) in hM3Dq animals (13.0±1.4 vs. 4.0±0.7) points towards higher levels of defensive responses (*U*_n1=56.5, n2=174.5_=1.5, *p*=0.0002). In addition, also the duration of SAPs was increased (8.9±2.4 s vs. 2.0±0.6 s) within the first 5 minutes of the EPM (hM3Dq×time: *F*_2,38_=3.59, *p*=0.0375, 2-way rmANOVA). Together, these results show that elevation of neuronal activity within the SC increased open-arm avoidance and risk assessment behavior even in blind LAB animals. Moreover, the pharmacogenetic stimulation of the SC could restore in part, the deficits in defensive responses to tactile stimuli.

### Reversing the High-anxiety Phenotype of HAB Mice

Similar to LAB mice, we used the MEMRI approach to identify the neural circuitry which potentially underlies the maladaptive defensive response pattern and increased open-arm avoidance behavior in HAB mice. A prominent brain structure found to exhibit increased manganese accumulation, was the ventrolateral, lateral (l) and dorsolateral (dl) periaqueductal gray (Fig 5A; for the complete MEMRI data set *see*). In order to assess the functional relationship of the PAG in the generation of the HAB behavioral phenotype, we implanted guide cannulae targeting the dl/lPAG (ML ±0.6 mm, AP −4.25 mm, DV −2.45 mm, needle protruded 500 μm) and injected 53.24 ng/100 nl (per hemisphere) of fluorescently labeled muscimol (MUSC), a potent GABA_A_-agonist (45 minutes before each experiment) which is comparable to 10 ng in 100 nl of ordinary muscimol. An exemplary image depicting the muscimol diffusion is shown in Fig 5B. The extent of muscimol diffusion of all animals (*N*=11) is shown in A) and comprised besides the lPAG, the dlPAG and partly the deep and intermediate layers of the SC. In order to test whether increased GABAergic signaling within the lPAG changes the extreme open-arm avoidance behavior of HAB mice, we have tested vehicle (aCSF, *N*=6) or MUSC (*N*=5) treated HAB mice (one VHC and two MUSC animals have been excluded from analysis due to deficient infusion) for their behavior on the EPM (Fig 5C-H). While only 20% of VHC treated animals accessed the open arm, all MUSC animals readily did so (Fisher’s exact, *p*=0.0152; Fig 5C+D). Further, MUSC treated animals spent significantly (*U*_n1=16, n2=50_=1.000, *p*=0.0116) more time on the open arm (44.4±9.4 % vs. 3.2±3.2 %; Fig 5E), less time in the closed arm (45.0±9.3 % vs. 88.1 ±5.8 %, *U*_n1=44, n2=22_=1.000, *p*=0.0087; Fig 5F) and showed increased locomotion (17.6±2.2 m vs. 8.0±1.2 m, *U*_n1=15, n2=61_=0.0, *p*=0.0043; Fig 5G). Time in center was unaffected (*see* B). These observations indicate a decrease in open-arm avoidance, which however, could be confounded by the increased activity. Therefore a decrease of active risk assessment (shown earlier to be sensitive to systemic diazepam treatment; *see* Fig 1F) in MUSC treated HAB mice (Fig 5H), namely number of SAPs (MUSC 5.0±3.5 vs. VHC 19.6±3.4, *U*_n1=45, n2=21_=0.0, *p*=0.008) supports the decrease in defensive responses. Moreover, the duration of SAPs was significantly decreased within the first 5 minutes of the EPM task with significant group effect (*F*_19_=13.71, *p*=0.0049, 2-way rmANOVA; Fig 5H). Finally, all animals were tested two times on two consecutive days for their disposition to vocalize during a 5 min TST, using a crossover design. Half of the animals received either VHC or MUSC treatment, which was swapped at the following day. Whereas 86% of VHC treated HAB mice emitted at least one sonic call during a 5 min TST, none of the MUSC treated animals vocalized (Fisher’s exact, *p*<0.0001; Fig 5I). All calls were in the sonic range. Fig 5J (*upper panel*) shows the spectral analysis of sonic vocalizing HAB mice (two mice have been excluded due to their low disposition of only short calls). Evidently HAB mice vocalize at a dominant frequency of 4090 Hz with a strong 1^st^ harmonic at 8180 Hz. All recordings were carried out using a USV transducer and were scored online using the heterodyne headphone output, thereby we can exclude that MUSC treated animals vocalized in the ultrasonic range only. Fig 5J (*lower panel*) shows an exemplary sonic call with the dominant frequency in the 4 kHz range, and formant harmonics up to approx. 16 kHz. In some calls (white asterisks) we can see that these harmonics even range up to 40-50 kHz, however these signals do not resemble any typical rodent ultrasonic call. These results, indicate an increased tonic activation of the PAG in HAB mice, which precipitates as an exaggerated open-arm avoidance behavior accompanied by a strong disposition to emit sonic calls, which could be reverted by low doses of muscimol.

**Figure 4.**
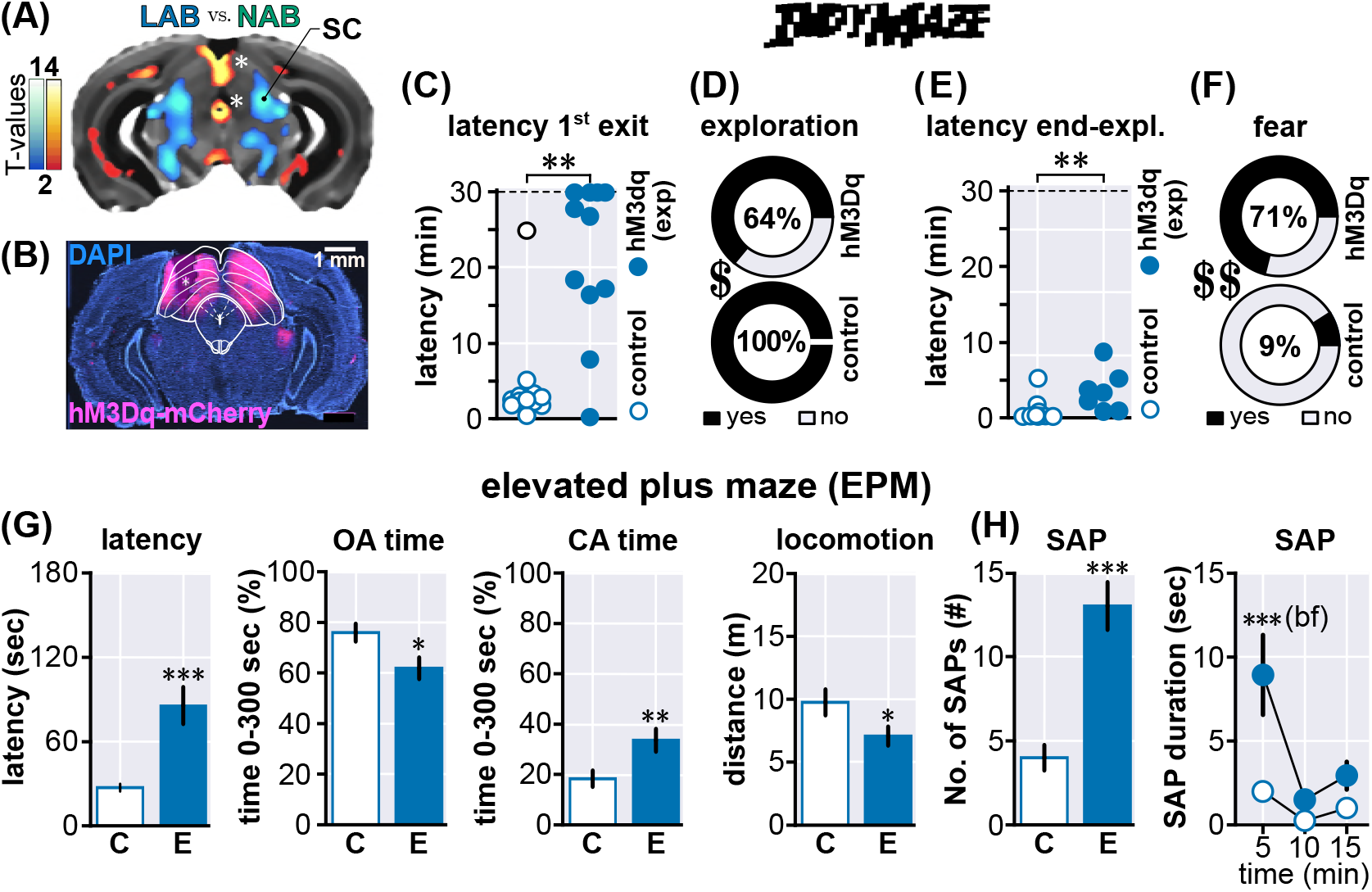
Reversing the Low-anxiety Phenotype of LAB Mice. (**A**) Manganese enhanced MRI (MEMRI) of LAB (*N*=30) vs. NAB (*N*=26) animals exhibited a significantly decreased accumulation of Mn^2+^ within the deep and intermediate layers of the superior colliculus of LAB. Warm colors indicate increased accumulation, cold colors indicate decreased accumulation in LAB as compared with NAB. The color brightness indicates the significance values. Asterisks mark signal artifacts within the aqueduct and above the superior colliculus due to line differences in brain templates. (**B**) Exemplary brain section at approximately the same slice location as the MEMRI data, depicting extent of viral expression (*magenta*) at the level of the superior colliculus. Shown in *cyan* is the nuclear 4’,6-diamidino-2-phenylindole (DAPI) counterstain. Overlaid are the outlines of the SC and PAG. Asterisk marks tissue lesion which occurred during sectioning. The effect of hM3Dq activation within the SC was studied using the IndyMaze task. Shown is the latency to first exit (**C**), the percentage of animals which explored the arena at all (**D**), latency to end-exploration (**E**) and the percentage of animals which showed a fear response to the ball (**F**). All animals were treated with 1 mg/kg clozapine-*N*-oxide (CNO) 45 minutes before the test. (**G**) In addition the same animals were tested for the behavior on the EPM (30 minutes), and latency to emerge, open arm (OA) time, closed-arm (CA) time and locomotion was assessed within the first 5 minutes. (H) Moreover the active risk assessment parameters, i.e. the total number of stretched-attend postures (SAP) and the duration of SAPs overtime (0-15 min) were scored. Asterisks indicate significance values obtained by Mann-Whitney test if not stated otherwise, * *p*<0.05, ** *p*<0.01, *** *p*<0.001;dollar signs indicate significance values obtained by Fisher’s exact tests, $ *p*<0.05, $$ *p*<0.01. Significance values obtained by 2-way rmANOVA, followed by Bonferroni post-hoc test are indicated with *bf*. Values are given as mean±SEM.

**Figure 5.**
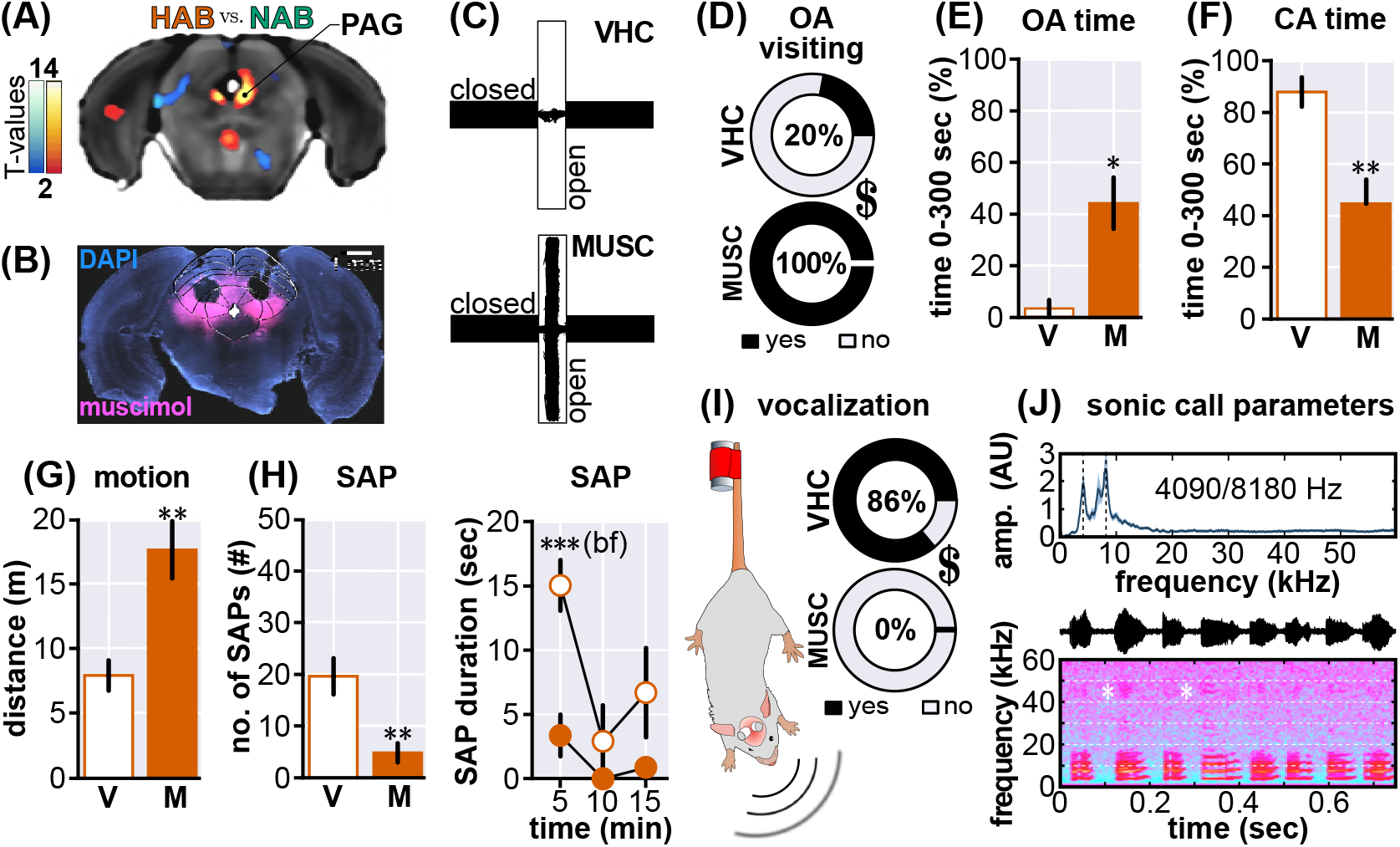
Reversing High-anxiety Phenotype of HAB Mice. (**A**) Manganese enhanced MRI (MEMRI) of HAB (*N*=31) vs. NAB (*N*=26) animals showed a significantly increased accumulation of Mn^2+^ within the periaqueductal gray of HAB. (**B**) Exemplary brain section at approximately the same slice location as the MEMRI image, depicting extent of fluorescently labeled muscimol (MUSC) diffusion (*magenta*) at the level of the periaqueductal gray. The nuclear DAPI counterstain is shown in *cyan* and overlaid by the outlines of the SC and PAG. Asterisk marks tissue lesion due to cannula placement. (**C**) Exemplary movement trace of a vehicle (VHC) and MUSC treated HAB mouse on the EPM. VHC (*N*=5) or MUSC (*N*=6) treated HAB animals were tested for their behavior on the EPM (30 minutes), and the percentage of open arm (OA) visiting animals (**D**), OA time (**E**), closed-arm (CA) time (**F**) and locomotion (**G**) was assessed within the first 5 minutes. (**H**) Moreover the active risk assessment parameters, i.e. the total number of stretched-attend postures (SAP) and the duration of SAPs overtime (0-15 min) were scored. (**I**) Finally, animals (*N*=14) have been treated with VHC and MUSC in a crossover design and subjected to a 5 min tail-suspension test (see cartoon), in order to assay the disposition to vocalize. (**J**) *Upper panel*: Spectral analysis of vocal call emitted by HAB (*N*=10). Depicted is the average (black) together with the SEM (blue). Dashed horizontal lines indicate the dominant frequency at 4090 Hz and the first harmonic at 8190 Hz. *Mid panel:* Exemplary call of a HAB animal;hull curve of raw signal. *Lower panel:* Sonogram of the same call. Note the formant structure of the harmonics. Asterisks denote rare and slight ultrasound artifacts within the 40-50 kHz range, which occur due to the expelled air itself. Asterisks indicate significance values obtained by Mann-Whitney test if not stated otherwise, * *p*<0.05, ** *p*<0.01, *** *p*<0.001;dollar signs indicate significance values obtained by Fisher’s exact tests, $ *p*<0.05, $$ *p*<0.01. Significance values obtained by 2-way rmANOVA, followed by Bonferroni post-hoc test are indicated with *bf*. Values are given as mean±SEM.

## Discussion

The inner feelings during states of anxiety, fear and panic of laboratory animals are not accessible to the experimenter. Instead one has to rely on behavioral and physiological readouts. Due to the rather continuous nature of the behavioral expression of anxiety, fear and panic, these states appear as a function of the animals defensive survival circuits. While this does not preclude a classification of the observed measures, it eliminates the possibility to ascribe a certain inner state to a behavioral category. In the description of the results presented in this study, we have limited ourselves to the use of *avoidance* for situations where the animal controls its exposure to a threat (emergence tasks), *active risk assessment* for the display of stretched-attend postures and *defensive responses* for directed and undirected flight and tonic immobility (freezing).

Previous studies investigating the neuropharmacological basis of altered open-arm avoidance of inbred mouse strains using the EPM, report a consistent dose-dependent *anxiolytic* effect of systemically administered benzodiazepines (*Rodgers et al., 1992; Cole and Rodgers, 1995; Holmes and Rodgers, 1999; Griebel et al., 2000*) which emphasizes the predictive validity of this testing situation. More recent studies which assess the involvement of specific brain areas in the expression and regulation of open-arm avoidance in rats and mice implicate the prefrontal cortex (PFC) (*Adhikari et al., 2010,2011; Kumar etal., 2013*), bed nucleus of the stria terminalis(BNST) (*Kim et al., 2013*), lateral septum (LS) to anterior hypothalamic area (AHA) projection (*Anthony et al., 2014*), medial septum (MS) (*Shin et al., 2009; Zhang et al., 2017*), septo-habenular pathway (*Yamaguchi et al., 2013*), basolateral amygdala (BLA) (*Sorregotti et al., 2018*) and specifically its projections towards the central nucleus of the amygdala (CeA) (*Tye et al., 2011*) and ventral hippocampus (vHPC) (*Felix-Ortiz et al., 2013*) and PFC (*Felix-Ortiz et al., 2016*), vHPC to PFC projection (*Padilla-Coreano et al., 2016*), vHPC-lateral hypothalamic area (LHA) projection (*Jimenez et al., 2018*), habenula (Hb) (*Pang et al., 2016*), interpeduncular nucleus (IPN) (*Zhao-Shea et al., 2015*), laterodorsal tegmentum (LdT) (*Yang et al., 2016*) but also the PAG (*Santos et al., 2003; Netto and Guimarães, 2004; Borelli and Brandão, 2008; Lima et al., 2008; Campos and Guimarães, 2009; Mendes-Gomes and Nunes-de Souza, 2009; Terzian et al., 2009; Muthuraju et al., 2016*) and the SC (*Muthuraju et al., 2016*). It is clear that all these brain structures cannot mediate the same types and aspects of avoidance behavior (e.g. social, olfactory, visual & auditory cues), but nevertheless they all modulate the same behavioral readout. While this broad spectrum of potentially involved circuits might be advantageous for initial behavioral screening purposes, the interpretation of the observed behavior on the EPM demands extra care. Consequently, referring to this behavior as *anxiety-like* is an oversimplification. In this study we presented data obtained from mouse lines which were generated under the simple assumption that the level of open-arm avoidance is a proxy for *anxiety (Krömer et al., 2005*).

It has to be noted that similar to the bidirectional selective breeding for open-arm avoidance in mice, there has been an earlier approach in rats wich resulted in high-anxiety and low-anxiety behaving animals (*for review see Landgraf and Wigger, 2003; Landgraf et al., 2007*). However, even though the findings obtained with HAB/LAB rats are highly relevant in the face of preclinical anxiety research, their discussion is beyond the scope of this study.

We have shown that the open-arm avoidance phenotypes of two mouse-lines which have been selectively bred for extremes in so-called *anxiety-like* behavior (HAB and LAB mice), based on their behavior on the EPM, are accompanied by paralleled changes in the level of defensive responses. This was demonstrated using two novel multi-sensory behavioral paradigms (Robocat, IndyMaze) which allowed the repeated assessment of innate escape behavior towards an approaching threatening stimulus. Further, we have discovered that LAB mice lack a functional retina due to a homozygous mutation in the *Pde6b*^rd1+/+^ allele which is indicative of the *retinal degeneration 1* (*rd1*) mutation and leads to blindness shortly after birth. Nonetheless, applying MEMRI-guided in-vivo neuronal circuit inquiry using the activating DREADD hM3Dq in the SC of LAB mice, we were able to demonstrate that increasing the neuronal activity within a midbrain multi-sensory integration circuit is sufficient to increase the level of innate defensive responses even in blind animals which ultimately precipitates as increased open-arm avoidance on the EPM. Further, using a similar approach but employing local injections of the potent GABA_A_-agonist muscimol into the PAG of HAB mice, we could show that also in this case a modulation of the neuronal activity within the midbrain survival circuits is sufficient to reverse the dominant open-arm avoidance phenotype.

### The Multidimensional Nature of Selective Breeding

Both bottom-up (which start from a defined genetic alteration or neuronal subpopulation or brain structure) and top-down approaches (which start from a distinct behavioral phenotype) hold the promise to decipher the molecular and cellular basis of anxiety-like behavior (*Anderzhanova et al., 2017*). The latter approach includes selective breeding and allows to study behavioral phenotypes on a polygenic background, which resembles the situation in most psychiatric diseases (*Landgraf et al., 2007; Anderzhanova et al., 2017*). It assumes that the resulting extremes in anxiety-like behavior reflect extremes in the normal distribution of the same behavioral trait (*Sartori et al., 2011b*). Accordingly, a direct comparison between the two extremes is expected to provide an optimal signal-to-noise ratio to disentangle molecular and cellular correlates of the phenotype. In fact, activity propagation through the amygdala circuit seems to support a dimensional shift from HAB via NAB to LAB phenotype (*Avrabos et al., 2013*), and many behavioral readouts show a similar pattern (*see* Table 1). The present study, however, demonstrates that this strategy might be misleading if not entirely wrong: First, measurements of differences in activity-dependent accumulation of Mn^2+^ did not reveal a single brain structure with bidirectional changes in signal intensity in HAB and LAB compared to NAB mice. Second, we identified impairments in sensory perception as a putative source of threat neglect in LAB mice. The *rd1* mutation freely segregates in many mouse strain populations, including CD1 (the ancestor strain used for the initial step of selective breeding). It stands for a nonsense mutation in the photoreceptor phosphodiesterase 6b (Pde6b). In case of homozygosity, the recessive mutation results in photoreceptor loss and retina degeneration. It is conceivable that the selective breeding of LAB is based, at least in part, on co-selection for *rd1* and the resulting physical blindness. Due to the lack of material, we cannot trace back the time point of first occurrence of homozygosity since the establishment of the LAB line more than 15 years ago (*Krömer et al., 2005*). In any case, data obtained in the past by direct comparison of LAB vs. HAB have to be (re-)interpreted with great care.

### In-*vivo* Imaging

We employed in vivo MEMRI imaging to investigate the neural basis of extremes in anxiety-like behavior. Other than expression of immediate early genes or accumulation of radioactive derivates of glucose which measure phasic changes in neuronal activity upon acute exposure to a threatening situation, repeated injections of MnCl_2_ are expected to result in intracerebral accumulation of Mn^2+^ also in cells with tonic (i.e., lasting) changes in neuronal activity. Importantly, MEMRI has the potential to non-invasively map whole-brain activity (*Bangasser DA, 2013; Bissig and Berkowitz, 2009; Chen et al., 2013, 2007; Eschenko et al., 2010; Hoch et al., 2013; Laine et al., 2017; Tang et al., 2016*). Mn^2+^ enters active neurons through voltage-gated calcium channels (*Drapeau and Nachshen, 1984*) (e.g. Ca_v_1.2; (*Bedenk et al., 2018)*), is transiently kept intracellularly (*Gavin et al., 1990*) and preferentially accumulates in projection terminals (*Bedenk et al., 2018*), which suggests its application for connectome analyses. Although our animals were not explicitly challenged, it has to be noted that the injection procedure per se may act as an acute stressor which triggers distinct neural responses in the different mouse lines. In fact, mice showed a prominent corticosterone secretion following treatment with Mn^2+^, which declined over the course of repeated injections (*Grünecker et al., 2010*).

Voxel-wise comparisons revealed a variety of brain structures with lower or higher Mn^2+^ accu-mulation in HAB or LAB vs. NAB. A detailed discussion of each of them is far beyond the scope of the present study. We wish to mention only a few prominent (and unexpected) findings such as the globus pallidus (GP) in LAB and the septo-hippocampal system in HAB. The GP is primarily associated with motor and associative functions (*Deniau et al., 2010*). However, it seems to play an important role also in the expression of aversive behaviour, including fear and anxiety (*Talalaenko et al., 2008*). Local administration of serotonin and glutamic acid into the GP effectively suppressed anxiety-like behaviour in the threatening situation avoidance test (*Talalaenko et al., 2008*), and; downregulation of the corticotropin releasing factor receptor 1 led to an anxiogenic effect (*Sztain-berget al., 2011*). Further clinical evidence for crucial involvement of the GP in anxiety mediated behaviour is given by the fact that deep brain stimulation within the GP was accompanied by a decrease of anxiety symptoms in depressive patients (*Kosel et al.,2007*) as well as patients suffering from Parkinson’s disease (*Tröster et al., 1997*). We cannot entirely rule out that the differences in GP activity may relate to line differences in general locomotor activity (*Yen et al., 2013,2015*). However, one would assume that increased motor activity, if at all, would lead to increased accumulation of Mn^2+^ in the motor network, resulting in an effect of order LAB > NAB not LAB < NAB.

Our finding of increased activation of the septal-hippocampal system in HAB is particularly interesting, given its suggested involvement in generalized anxiety disorder (*Gray and McNaughton, 2000*). Septal lesions were reported to increase the time spent on the open arm of the EPM and to decrease the time spent burying the shock probe (*Menard and Treit, 1996*), and the local administration of the arginine vasopressin receptor antagonist into the septum of rats led to an increased time spent on the open arm of the EPM (*Liebsch et al., 1996*). Septal neurons which express the corticotropin releasing factor receptor 2 project to the hypothalamus and promote anxiety-like ι behavior (*Anthony et al., 2014*). Also the hippocampus has been implicated in anxiety-like behavior (*Bannerman et al., 2004*), in particular its ventral part (*Felix-Ortiz et al., 2013; Padilla-Coreano et al., 2016; Jimenez et al., 2018*). Therefore, it is tempting to assume that the higher activity status of the I hippocampus in HAB mice indicated by increased Mn^2+^ accumulation causally links to exaggerated fear and anxiety-like behavior shown by the animals. This interpretation is supported by PET-studies; in rhesus monkey which report increased brain activity in the hippocampus of hyperanxious animals i (*Oler et al., 2010*). Interestingly, also LAB mice show higher Mn^2+^ accumulation in the hippocampus formation, which seems to contradict our interpretation. However, we observed a prominent signal; at the level of the dorsal dentate gyrus, which has been associated with hyperlocomotion and decreased anxiety before (*Kheirbek et al., 2013*), thus resembling the LAB phenotype (*Yen et al., 2013*).

### Midbrain Structures Control the Level of Open-arm Avoidance, Risk Assessment and Defensive Behavior

I Among the many brain structures with different accumulation of Mn^2+^, there were also parts of the midbrain/tectum, which showed reduced (e.g., superior and inferior colliculus in LAB mice) or; enhanced (PAG in HAB mice) signal intensities. The superior colliculus is a multimodal sensory-motor structure that receives inputs from the retina and somatosensory cortex (*King, 2004; Shi et al., 2017*). Efferences from the superior colliculus trigger a variety of defensive responses (*Shang et al., 2015; Evans et al., 2018*). Therefore, the reduced activity status of the superior colliculus may reflect both reduced sensory inputs (i.e., threat detection; (*Almada et al., 2018*)) and reduced threat responding. Physical blindness alone is insufficient to explain the behavioral phenotype in the IndyMaze (where LAB mice failed to show flight responses even after contact with the styrofoam ball) and on the EPM (low level of risk assessment). Indeed, we could reduce the ‘emotional blindness’ by chemogenetic activation of the superior colliculus. In HAB, the enhanced Mn^2+^accumulation spans the entire caudal part of the PAG and resembles the enhanced expression of c-Fos in HAB mice which had been confined to an open arm of the EPM (*Muigg et al., 2009*). The increased activity in ventral parts of the PAG correspond to the prevalence of HAB mice for showing passive defensive responses (*Bandler R, 2000; Tovote et al., 2015*)(*see also* Table 1). The increased activity in dorsal parts is more surprising, given their association with active defensive responses (*Bandler R, 2000*). Only recently we could demonstrate that HAB mice show exaggerated active fear to an approaching robo-beetle (*Heinz et al., 2017*), which is in accordance with the exaggerated flight responses to the Robocat shown in the present study. Remarkably, inactivation of the dorsal PAG led to the most striking changes in EPM behavior observed so far in this mouse line. This also applied to the reduction in risk assessement and the complete absence of defensive vocalization.

### Of Fear and Anxiety

In animals, anxious states are prototypically assessed in exploration- or interaction-based tasks which involve approach-avoidance conflicts (*Belzung and Griebel, 2001; Millan, 2003; Cryan and Holmes, 2005; Sousa et al., 2006*). Avoidance measures alone are insufficient to describe the behavioral phenotype, since they might be confounded by alterations in exploratory drive. Therefore, it is strongly recommended to additionally assess ethobehavioral parameters which reflect approach behavior torwards a (potentially) threatening environment/object (*Gray and McNaughton, 2000*). Here we report bidirectional changes in risk assessment (*McNaughton and Corr, 2018*) on the EPM in HAB and LAB vs. NAB mice, which were sensitive to diazepam treatment and could be reverted by pharmacological or chemogenetic interventions at midbrain/tectal structures. Open-arm exploration, in contrast, was insensitive to diazepam and, thus, seems to be less suited as a measure of anxiety-like behavior. This might be ascribed to the stringent selection process over the generations (*Krömer et al., 2005*), and the threat intensity due to the combination of height and open spaces. Accordingly, during the first phase of the IndyMaze, when mice have to leave the home compartment to explore the hollow way engulfed by side and end arms, HAB mice showed a strong increase in emergence latencies. This time, however, this parameter was sensitive to diazepam treatment, possibly because of a less threatening impact of the test situation. Importantly, HAB mice readily explored the entire setup, once they had left the home compartment. Therefore, differences in EPM or IndyMaze exploration cannot be ascribed to a general lack in motivation/exploratory drive or locomotor behavior, but to state anxiety. To study the consequences of extremes in anxiety-like behavior on defensive responses to explicit threat, we decided to develop ethobehavioral tasks (*Pellman and Kim, 2016*), which allow for the measurements of active defensive responses as a function of defensive distance and to judge their adaptive vs. maladaptive nature. In the IndyMaze, mice were confronted with an approaching styrofoam ball, which spans the entire width of the hollow way. Whereas both HAB and NAB escaped from the ball, the behavior of LAB mice was clearly maldadaptive, since virtually all mice were overrun by the ball at least once (without physical harm). As discussed before, increased neuronal activity in the superior colliculus reestablished flight behavior in the majority of the animals, demonstrating that not exclusively sensory deficits (i.e, physical blindness) can explain the deficits in defensive behavior. A similar picture emerged from the Robocat exposure, which resembles the robogator described before (*Choi and Kim, 2010; Amir et al., 2015; Pare and Quirk, 2017*). Again, LAB mice were at high risk to collide with the Robocat. This time, also the behavior of HAB mice turned out to be maldadaptive, since no mouse could bypass the Robocat even if not at risk to collide with it. Together with our previous observation of increased avoidance of an approaching robo-beetle (*Heinz et al., 2017*), this finding suggests that selection for high levels of anxiety-like behavior on the EPM coincides with exaggerated defensive responses, both passive (*see* Table 1) and active.

## Conclusion

Using well-established mouse lines with extremes in anxiety-like behavior, we demonstrate that extremes in anxiety coincide with (i) extremes in defensive responses to an approaching threat and (ii) tonic changes in neuronal activity, among others in midbrain/tectal structures, (iii) which – if reverted – ameliorated both fear- and anxiety-like behavior. In addition, we provide evidence for (iv) the multidimensional nature of increased vs. decreased defensive behavior, which may include deficits in sensory perception. Our results challenge the uncritical use of the anthropomorphic terms anxiety or anxiety-like for the description of mouse behavior on the EPM or in other exploration-based tasks (*for review see Ennaceur, 2014*), as they imply higher cognitive processes, which are not necessarily in place. The explicit fear of height (acrophobia) and/or open spaces (agoraphobia) sufficiently explains the lack of open-arm exploration. The recently initiated discussion about the uncritical use of the term ‘fear’ where ‘threat’ would be more appropriate (*LeDoux, 2012*) has forced the scientific community to reconsider its terminology, even though the term ‘fear’ still keeps its merits (*LeDoux, 2014, 2017*). We face a similar if not more eminent problem, if we uncritically use the term ‘anxiety’ in translational studies on animal behavior. Instead, we should describe defensive responses as they are, preferentially along the continuum of the *predatory imminence model (Perusini and Fanselow, 2015*).

## Methods and Materials

### Animals

Adult (3-8 months), male mice of the following strains have been used: C57BL/6N (B6) (*N*=12), HAB (*N*=154), NAB (*N*=76), LAB (*N*=99), CD1 (*N*=12), resulting in a total number of 353 animals. All animals were bred in the animal facilities of the Max Planck Institute of Biochemistry, Martinsried, Germany. The animals were group-housed (2-4 animals per cage) under standard housing conditions: 12h/12h inverted light-dark cycle (light off at 8AM), temperature 23±1 °C, food and water *ad libitum*. Experimental procedures were approved (55.2-1-54-2531: 44-09,188-12,142-12, 133-06, 08-16) by the State of Bavaria (Regierung von Oberbayern, Munich, Germany). Animal husbandry and experiments were performed in strict compliance with the European Economic Community (EEC) recommendations for the care and use of laboratory animals (2010/63/EU). All efforts were done to minimize the number of experimental subjects and to preclude any animal suffering.

### Drugs

The anxiolytic diazepam (Diazepam-Lipuro^®^, Braun Melsungen, Germany) was diluted in physiological saline (vehicle) and injected systemically (1 mg/kg, i.p.) using a volume of 100 *μ*l per 10 g body weight. Clozapine-*N*-oxide (CNO, Tocris #4936) was dissolved in dimethyl sulfoxide (DMSO, Sigma-Aldrich, #472301) at a stock concentration of 75 mM and stored at −20°C. The final concentration of CNO was 292 *μ*M (1 mg/kg at 100 *μ*l per 10 g body weight) in saline (<0.5% DMSO). Muscimol MUSC (Sigma-Aldrich, #M1523) and fluorescently-labeled muscimol (fMUSC) (Bodipy^®^TMR-X conj. Thermo Fisher Sc. M23400) was dissolved in artificial cerebrospinal fluid (aCSF) (*Baarendse et al., 2008*). MUSC itself has a molecular weight of 114.1 g/mol whereas the fMUSC (MW 607.46 g/mol) is 5.324× heavier. In previous experiments with MUSC, we found a concentration of 10 ng/100 nl (876.4 *μ*M) most effective, therefore we have used 53.24 ng/100 nl fMUSC to achieve the same physiological effect. As the fMUSC is poorly water-soluble, we dissolved 1 mg in 1.878 ml aCSF to reach a final ready to use concentration of 876.6 *μ*M. Whereas the EPM experiments were conducted using MUSC, the vocalization experiments only involved fMUSC. The vocalization experiment was carried out using a crossover design: half of the animals received fMUSC on the first day, whereas the other half received VHC (aCSF). On the next day the treatment was switched. 1-3h after the experiment the animals which received fMUSC were transcardially perfused (4% PFA in PBS), whereas the remaining animals received another injection of fMUSC on the following day and were also perfused 1-3h after the injection.

### Behavioral Tests

#### Elevated Plus Maze

The elevated plus maze (EPM) apparatus consisted of two open (L30×W5 cm) and two closed (L30×W5×H15 cm) arms which were connected via a central platform (L5×W5 cm). All parts of the EPM were made of dark gray PVC. The apparatus was elevated 37 cm above a table (H50 cm), which was placed in the center of the dim illuminated experimental room (5×4 m). The light intensity (luminous flux) at the open arms was 7 lux. Before the experiment every subject was tested for the disposition to emit sonic vocalizations by lifting them 3 times from the grid cage top (*Whitney, 1970*). At the beginning of each trial, the animal was placed near the central platform facing a closed arm. Each trial lasted for 30 minutes and was videotaped. The animals behavior was analyzed using a behavioral tracking software (ANY-maze, Stoelting CO., USA) and the percentage of time spent on the open (OA) and closed (CA) arm and the central zone (time in center) as well as the total distance traveled were determined. In order to render these results comparable to other EPM experiments found in the literature, the data (except for latency and stretched-attend postures, SAPs) of the first 5 minutes of each trials is reported. Other behavioral parameters which were analyzed by an experienced observer, blind to the experimental conditions, included number and duration of SAPs within the first 15 minutes of each trial, and the latency for the first full entry to the open arm (all four paws) within the entire 30 minutes exposure. After the trial the fecal boli on the EPM apparatus were counted as a measure of autonomic arousal. In between the trials the apparatus was cleaned with tap water containing detergent, and was subsequently dried with tissues.

#### Robocat Task

The Robocat (for a detailed explanation of the task *see*) has been inspired by the Robogator (*Choi and Kim, 2010*). It is a four-wheeled robot (Lego Mindstorms), equipped with ultrasound range finders and programmed to advance for 25 cm (speed 25 cm/s) once a movement has been detected within the sensor range of 50 cm. Despite the name suggests, no extra effort has been invested to disguise the robot as a cat, except two little cardboard ears. The task is conducted within a longitudinal arena (H35×W50×L150 cm, whereby the robot is placed 125 cm away from the start compartment (H35×W50×L12.5). The access to the arena is provided via a sliding door, operated by the experimenter, and the natural exploratory drive (neither bait, nor food or water deprivation used) ultimately leads to the mouse-robot encounter. Once the mouse triggers the robot, its movements typically evoke a robust flight response and the mouse retrieves to the start compartment. All animals were first pre-exposed to the entire setup with unrestricted access to the arena (sliding door opened) in absence of the Robocat. On the following consecutive 3 days each animal was subjected to habituation trials which consisted of 10 minutes acclimatization within the start compartment (to enable the mice to form a home base), followed by 10 minutes of free exploration in the arena, again without the Robocat. The test trial on day 4 was conducted in identical manner, except that the Robocat was placed in the arena. During the test trial, the animals typically activated the Robocat several times. All trials were videotaped and the behavior was analyzed offline by an experienced observer, blind to the experimental conditions. The behavioral readouts were flight (activation + retrieval), bypass (activation but tolerance to the approaching Robocat which is bypassed by the animals) or collision, and were counted if observed at least once. Only animals which activated the Robocat at least once were considered for analysis.

#### IndyMaze Task

Inspired by the movie *Indiana Jones and the Raiders of the Lost Ark* (*Spielberg and Marshall, 1981*), the IndyMaze is conducted within a narrow, stretched arena (H35×W16×L150 cm), which was divided into six equidistant (25 cm) sectors. To one end of the arena, a small custom-made plexiglass cage (H30×W16×L25 cm), equipped with bedding material, was connected which served as a home compartment. The arena itself was slightly tilted towards the home compartment and indirectly lit (<10 lux). To enter the arena, the animals had to climb over a small barrier (height: 2 cm). This prevented the animals from ‘accidentally’ dropping into the arena and forced them to explicitly decide when to initiate its exploration. For the task, each animal was first placed into the home compartment and was allowed for a maximal duration of 30 minutes to step (with four paws) into the arena (latency 1^st^ entrance). Once the animal had entered the arena, the time to reach the last sector was noted (latency for end-exploration). After the end exploration, the animals typically retrieved to the home compartment or were gently forced to do so by the experimenter. With low latency the animals re-entered the arena but this time a styrofoam ball (Ø15cm, 100 g) was introduced at the last sector, which was allowed to roll (25 cm/s) towards the animal once it passed the midline (75 cm). The animals either responded with (a) preemptive flight or a retrieved once the ball hit them (both counted as defensive responses), or (b) they were overrun by the ball (without any physical harm) and continued to explore the arena. The threat exposure part of the behavioral paradigm was carried out for a maximal duration of 30 minutes or once the animals had encountered the ball three times. The behavior was scored online by the experimenter during the task unaware of the mouse line.

#### Optomotor Response

In order to assess the visual performance of male C57BL/6N, CD1, HAB, NAB and LAB mice (*N*=12), the animals’ optomotor response (*Abdeljalil et al., 2005*) has been tested using the rotating drum task. The task is based on the mouses’ predisposition to fixate on moving vertical black/white stripes and follow their rotation with short movement bouts, involving the entire head. By decreasing the stripe width, higher visual acuity is necessary to resolve the stripes. The apparatus consisted of a rotating cylinder (drum, Ø33 cm, height 35 cm), whose inner walls were lined with an alternating black/white stripe pattern using a stripe width of 2.88 cm, giving a spatial frequency of 0.05 cycles per degree (cyc/deg, *r*=16.5 cm, arc length per black/white cycle 5.76, angle 20°). During the task, the animals were placed within the center of the drum on a Ø11.5 cm fan grid which was mounted 16 cm above the bottom. The rotation of the drum was controlled via a custom-built microprocessor-based motor driving circuit which operated a geared motor. The rotational speed of the drum was set to 2.5 rounds per minute (rpm). For the task the animals were placed into the drum for 1 minute to acclimatize (bright illumination 500 lux) and subsequently the drum started to rotate for 60 seconds clockwise, followed by a 30 seconds break and then rotated in counter-clockwise direction for additional 60 seconds. All experiments were videotaped and analyzed offline (blind to the strains with same fur color, i.e. CD1, HAB, NAB and LAB), whereas every head movement was scored as an optomotor response if it was directed into the same rotational direction as the drum. This modified version of the original task *[Abdeljalil et al., 2005*) does certainly not allow to make detailed statements regarding different levels of visual acuity, though it is sufficient to assess the general visual performance of the mouse strains in question.

### Physiological Measurements

#### Electroretinography

In order to assess the retinal function of male C57BL/6N, CD1, HAB, NAB and LAB animals (*N*=6 each), flash-evoked electroretinographic (fERG) measures in the anesthetized animals have been employed. To this end, the animals were dark-adapted for > 3 h prior to the experiment. Under dim red light (650 nm) illumination, the animals were weighed and received analgesic treatment (200 mg/kg Novalgin/Metamizol s.c. in saline in a concentration to obtain 100 *μ*|/10 g of body weight) and subsequently transferred from their home-cage to the anesthesia chamber (isoflurane 4%). After reaching surgical tolerance, indicated by the absence of the eye-lid and paw-withdrawal reflex, the animals were transferred to a modified stereotaxic frame were the anesthesia was maintained with isoflurane (2-3 % in oxygenated air, using an oxygen concentrator, EverFlo). The body temperature was monitored and controlled (37.5°C) using an animal temperature controller (WPI Inc. #ATC2000) in combination with a small rodent rectal temperature probe (WPI Inc. #RET-3) and a small heating-pad (15×10 cm) with built-in RTD sensor (WPI Inc. #61830) with an additional silicone pad to ensure maximal heat transfer (WPI Inc. #503573). For the analgesic treatment to have an effect, the animals was allowed to reach a stable anesthesia for >15 min, while the eyes were kept moisturized with 0.9 % (w/v) physiological sodium chloride solution (saline). subsequently the pupils were dilated maximally using 2.5 % phenylephrine (Sigma #P6126, in PBS, pH adjusted to 7.0) and 1 % (w/v) atropine (Sigma # A0132, in PBS, pH adjusted to 7.0) and the eyes were henceforward kept moisturized using 1 % methyl cellulose (Carl Roth #8421) in saline. The ERG electrodes were custom made using Ø200 μm uncoated gold wire wound to form Ø3 mm loops and were placed gently on the eyes of the animal. A stainless steel wire wrapped around the animal’s tail served as the ground electrode. All signals were bandpass filtered at 0.1-300 Hz and sampled at 30 kHz using the Open-Ephys (*Siegle et al., 2017*) system in conjunction with a headstage based on the Intan RHD2132 integrated extracellular amplifier circuit. The animals left eye was covered with a piece of light proof black PVC and additionally shielded from the right side using aluminum foil. The animals right side was stimulated using a Ping-Pong ball which was cut in half (*Green et al., 1997*) and illuminated with a white LED (Osram Oslon LUW CN7N) which was controlled via a custom-made constant current source. Thereby scotopic and photopic (3 lux background illumination) measurements were carried out which involved the display of 32 light flashes (per condition) of 40-180 *μ*s length at a frequency of 1 Hz at three different light intensities. The light intensities were measured (65 lux, 225 lux, 420 lux) using a hand-held lux meter (Iso-Tech ILM 1335) and the respective log Φ·rod^-1^ s^-1^ values were calculated using the following relation:

1 photopic lux = 650 photoisomerizations (Φ)·rod^-1^ ·s^-1^ (*Pugh et al., 1998*).

All 32 acquired responses per condition were averaged and the datasets were further analyzed using custom Python2.7 scripts.

#### Vocalizations

During the normal animal care taking procedures, it was realized that HAB mice have a strong disposition to vocalize in the audible hearing range, if lifted at their tails (e.g. at changing cages) and especially when they lose grip from a grid cage top. Although there have been previous attempts to standardize this cage-grid vocalization test (*Whitney, 1970*), in our study the tail-suspension test (TST) was employed, a behavioral test which typically aims to assess depression-like behavior in mice (*Steru et al., 1985*). For this test, the animal was affixed roughly 2 cm above the tail root to a Ø5 mm vertical stainless steel rod (20 cm above ground) using heat sterilization tape. Other tapes can be used, but it was found that this sort of material is characterized by its rather low adhesion to murine skin and its excellent removability without introducing skin irritations. The test was carried out within a sound-attenuating chamber. The test duration was 5 minutes, and the animals vocalization was monitored using high-quality sonic/ultrasonic recording equipment (Avisoft UltraSoundGate USG 116-200, condenser microphone CM16/CMPA). Offline analysis was carried out using custom written Python2.7 scripts.

### Standard Laboratory Procedures & Analysis

#### Stereotaxic Implantation and Virus Injections

All stereotaxic surgical procedures were carried out similarly and shall be briefly described. Specifics for cannulae implantations and virus injections are provided if necessary. Before the surgery, the animal was weighed and analgesic treatment (200 mg/kg Vetalgin, Intervet, in saline, s.c.) was administered 15 minutes prior to any other interventions. During this time, all surgical instruments have been heat sterilized and wiped with 70 % ethanol. Than the animal was transferred to the anesthesia induction chamber and slowly anesthetized with isoflurane (0-4 % in oxygenated air, EverFlo Oxygen Concentrator). The absence of the eyelid and paw withdrawal reflex indicated surgical tolerance and the animal was transferred to the stereotaxic frame (Leica Biosystems, AngleTwo), where it was fixed using non-rupture/non-traumatic ear bars and a snout clamp. The anesthesia was kept constant with 2-2.5 % isoflurane, while the animals body temperature was constantly monitored and controlled (37.5°C) using a rodent rectal probe, heating blanket and a animal temperature controller (WPI Inc. ATC2000). The eyes were kept moisturized using eye ointment (Bepanthen^®^ eye and nose ointment). Further the animals head was shaved using either serrated scissors or an electric shaver. Excess cut hair was removed with cotton swabs soaked with lidocaine (Sigma #L7757,10 % (w/v) in 70 % ethanol) which in addition exerted an additional cutaneous analgesic effect. Using sharp scissor, the skin above the skull was opened from 1 mm caudal to lambda to 2 mm rostral to bregma. The periosteum was removed with clean cotton swabs soaked in lidocaine solution followed by 3 % hydrogen peroxide. Now, using a small and stiff probe the AngleTwo system was calibrated with the position of bregma and lambda and medial-lateral (ML) and dorsoventral (DV) deviations were corrected if necessary to read less than 50 *μ*m utilizing the manufacturing tolerances of the mouse skull adapters’ dovetail rails. In order to correct the skull rotation, two contra-lateral coordinates on the skull surface were targeted (ML ±2.0 mm, AP −1.82 mm) and the respective DV coordinates were noted. If a deviation >50 *μ*m was noticed, the ear bars were released and the initial rotation was corrected. Once the position of the skull was sufficiently accurate, implantation or virus injection was conducted. After these procedures the animals were weighed and their general health and healing status was assessed and recorded on a daily basis for 5 consecutive days and in addition the animals received post-surgical analgesic treatment (1 mg/kg Metacam, Böhringer Ingelheim, in saline, s.c., daily). For viral injections, a 5 *μ*| Hamilton syringe (7634-01/00) equipped with a blunt 33 gauge needle or a 10 *μ*| WPI Inc. syringe (NANOFIL) equipped with a 34 gauge beveled needle (NF34BV-2) in conjunction with a motorized micropump (WPI Inc. UMP3) and the respective micropump controller (WPI Inc. MICRO4) was used. The injection rate was set to 80 nl/min. For the experiments involving the pharmacogenetic manipulation of the SC in LAB mice, 350 nl of adeno-associated-virus serotype-5 (AAV), expressing either the active DREADD (AAV5-CaMKII*α*-hM3Dq-mCherry, #AV6333, *N*=12) or just the reporter fluorophore (controls, AAV5-CaMKII*α*-mCherry, #AV4809c, *N*=12), have been injected (ML ±0.9 mm, AP −3.64 mm, DV −1.75 mm). All viruses were purchased from the Gene Therapy Center Vector Core of the University of North Carolina, Chapel Hill and were diluted, using 350 mM NaCl solution, to reach a target titer of 1.7×10^12^ vg/ml. For the injection, first, the target drilling site was marked with a pencil on the skull surface, and the skull was penetrated using a Ø0.5 mm burr with counterclockwise concentric movements until the intact dura mater becomes visible. Using a hypodermic needle, whose foremost sharp tip was gently bent to the outside by tipping it onto a polished stainless steel surface in order to form a micro-miniature hook-like instrument, was used to first remove the remaining skull pieces and secondly to open the dura at the site of injection. The injection needle was slowly lowered to reach the target site and the injection was initiated. After the injection the needle was raised for 100 μm and left for additional 10 minutes in order to allow the virus to diffuse. Subsequently, the needle was removed and the procedure was repeated on the contralateral side. During the injection the wound was kept moisturized using saline, in order to prevent brain tissue from sticking onto the needle and to aid the subsequent cutaneous suture. After the injection, using resorbable, sterile, surgical needled suture material (VetSuture fastPGLA 5/0, 13 mm reverse cutting needle 3/8), the wound was closed with 4-6 intermittent stitches, and treated with iodine solution (Braunol^®^). The incubation time for the virus to reach stable expression was >5 weeks.

#### Guide Cannula Implantation & Local Muscimol Injections

For the local injection of muscimol within the lPAG of HAB mice (*N*=14), two 3.0 mm long, 26 gauge guide cannulae (WPI Inc.) have been implanted using an angle of ±25° at ML ±1.02 mm, AP −4.25 mm and DV-1.55 mm. As the internal injection needle had a length of 4.0 mm, the ultimate injection site was ML ±0.6 mm, AP −4.25 mm and DV −2.45 mm. One skull screw per hemisphere above the hippocampus (ML ±1.5, AP −1.27) allowed a mechanically stable attachment of the cannulae to the skull using dental cement (Paladur^®^, Heraeus-Kulzer). Iodine solution (Braunol^®^) was used to disinfect the wound. After the implantation, dummy injection needles with a dust cap and a length of 3.5 mm were inserted into the guide cannulae in order to prevent clogging. The animals were allowed to recover for more than 2 weeks after the surgery. The injection of MUSC or fMUSC or vehicle (aCSF) before the EPM and vocalization task was conducted in the anesthetized (2-2.5 % isoflurane) animal. The injection was carried out using an ultra micropump (WPI Inc. UMP3) and the injection rate was set to 100 nl/min whereby volume of 100 nl was injected. 45 minutes after the injection, the animals were subjected to the behavioral paradigm.

#### Histology

For histological verification of injection and implantation sites, the animals were deeply anesthetized using a mixture of ketamine (50 mg/kg, Essex Pharma GmbH, Germany) and xylazinhydrochloride (5 mg/kg, Rompun, Bayer Health Care, Germany) injected systemically (100 *μ*| per 10 g body weight, i.p.). subsequently the animals were given an overdose of isoflurane to induce respiratory arrest (final anesthesia) and transcardially perfused with cold physiological saline followed by 4% (w/v) paraformaldehyde (PFA) in phosphate buffered saline (PBS, final concentrations in mM: 136.89 NaCl, 2.68 KCl, 10 Na_2_HPO_4_,1.76 KH_2_PO_4_; pH adjusted to 7.4 using HCl). The brains of the animals were post-fixed in PFA solution for >24 h at 4°C. In order to prevent the implant tracks from collapsing upon removal, the entire heads of the animals were post-fixed for >48 h. The brains were further placed in 30% (w/v) sucrose in PBS solution for >36 h at 4°C for cryoprotection in order to increase tissue rigidity. subsequently the brains were dry dabbed and carefully frozen by repeatedly dipping the brain, held at the medulla, into the cold 2-methylbutane on dry ice and stored at −80°C. Coronal tissue sections of 35 *μ*m, cut in several series, were prepared using a cryostat (Thermo Scientific Microm HM560). Sections were collected directly on microscopy slides (SuperFrost^®^, Menzel-Gläser, Germany). For proteinaceous fluorophores the specimens were covered and preserved using antifade mounting medium (VECTASHIELD^®^ HardSet H-1500, VECTOR Laboratories, UK) containing the nuclear counterstain 4’,6-diamidino-2-phenylindole (DAPI). Some series were stained using the standard Nissl staining method in order to reveal the gross anatomical structures. In brief, the specimens were dehydrated using (in v/v) 80%, 90%, 2×100% ethanol (30 seconds per step), stained in 0.1% (w/v) cresyl violet solution in double distilled water acidified with 300 *μ*| glacial acetic acid for 30 seconds. subsequently the specimens were differentiated in 100% isopropyl alcohol (for 30 seconds)followed by 100% xylene for (>5 min). The cresyl violet stained sections were covered and preserved using DPX mounting medium. For the preparation of retinal section the eyes of the perfused animals were removed and stored in 4% PFA at 4°C and the retinas were extracted. Retinal sections (30 *μ*m) were obtained (*Ivanova et al., 2013*) using a cryostat and the specimens were stained with haematoxylin and eosin.

#### Genotyping for *Pde6b*^rd1^

The genotyping for *Pde6b*^rd1^ was carried out according to Chang et al. 2013 (*Chang et al., 2013*). In brief, genomic DNA was extracted from tail biopsies (B6, CD1, HAB, NAB, LAB, *N*=4 per strain) by adding 100 *μ*| 50 mM NaOH aqueous solution to each sample (per 1.5 mL reaction tube) followed by 30 minutes incubation at 99°C. Subsequently the samples were allowed to cool down and 30 *μ*| of 1 M Tris-HCl aqueous solution was added per sample. Finally the samples were thoroughly vortexed and cell debris was removed by brief centrifugation and the samples were stored at −20°C. For the polymerase chain reaction (PCR), 2.5 *μ*| PCR buffer (Thermo Scientific, ThermoPrimeTaq 10x Buffer), 2.5 *μ*| MgCl_2_ (25 mM), 1 *μ*| deoxynucleoside triphosphate (dNTP, 10 mM) mix (Thermo Scientific, 18427-088), 1 *μ*| dissolved G1 primer, 1 *μ*| G2 primer, 1 *μ*| XMV primer, 0.2 *μ Taq* DNA polymerase (Thermo Scientific, ThermoPrime, #AB-0301/B) and 14.8 *μ*| double distilled water was mixed with 1 *μ*| of genomic DNA solution. The primer sequences were as follows: G1 (5′-CCTGCATGTGAACCCAGTATTCT ATC-3′), G2 (5′-CTACAGCCCCTCTCCAAGGTTTATAG-3′) and XMV (5′-AAGCTA GCTGCAGTAACGCCATTT-3′). The idea of this three primer design is that while G1 and G2 result in a PCR product of 240 base pairs (bp) from normal non-mutant animals, G2 and XMV generate a larger (560 bp) product from the *rd1* mutant allele. The thermal cycler PCR protocol consisted of the following steps: denaturation for 3 minutes at 95°C, followed by 34 cycles of annealing (30 seconds, 55°C) and extension (1 minute, 72°C) terminated with a final cycle at 72°C for 5 minutes and the subsequent incubation at 4°C. The amplified DNA was analyzed using agarose gel electrophoresis and a subsequent ethidium bromide staining.

#### Manganese-enhanced MRI

The animals (HAB *N*=31, NAB *N*=26, LAB *N*=30) were injected with a low dose of manganese chloride (30 mg/kg in saline, i.p.) for eight consecutive days (8×30/24 h) prior to the scanning procedure, see Grüenecker et al. 2010 (*Grünecker et al., 2010*). The MRI experiments were performed in a 7T MRI scanner (Avance Biospec 70/30, Bruker BioSpin, Ettlingen, Germany) at 24 h after the last injection, with the animals being anesthetized with isoflurane (≈ 1.5-1.7% in oxygenated air). Body temperature was monitored and kept constant in the range 34-36°C. A saddle-shaped receiver coil was used for signal acquisition. T_1_-weighted images were acquired using a 3D gradient echo pulse sequence (repetition time TR = 50 ms, echo time TE = 3.2 ms) using a matrix of 128×128×128 at a field of view of 16×16×18 mm^3^, yielding a final resolution of 125×125×140.6 *μ*m^3^. 10 averages were acquired. In addition, 3D T2-weighted images were acquired using a rapid acquisition relaxation enhanced (RARE) pulse sequence (TR = 1 s, TE = 10 ms) with the same spatial resolution as mentioned above, and two averages. This resulted in a total imaging time of approximately 2 hours per animal. The reconstructed images (Paravision, Bruker BioSpin, Ettlingen, Germany) were further analyzed using the statistical parametric mapping package SPM5 (using the spmmouse toolbox) and SPM8 (using the new segment option for bias correction) (www.fil.ion.ucl.ac.uk/spm/).

The acquired images of all animals were segmented exploiting mouse specific tissue probability maps, and bias corrected images were obtained. Then, images were spatially normalized in several steps: 1. Normalization of all images (including brain and extracranial tissue) to a representative single animal image and calculation of the mean normalized image. 2. Creation of a brain mask on the mean normalized image. Brain extraction in native space using the back-transformed mean brain mask. 3. Normalization of the brain extracted images to the group template. Finally, images were smoothed using a Gaussian kernel of eight times the image resolution. Data were further analyzed in SPM using a full factorial design with three conditions (HAB, NAB and LAB), global mean correction and global normalization using ANCOVA. A pairwise voxel-based comparison between HAB vs. NAB and LAB vs. NAB (FDR *p*<0.001, cluster extent >20) revealed the differential manganese accumulation ().

### Statistical Analysis

All data are presented as mean values ± standard error (SEM). Statistical analysis has been performed using GraphPad Prism 7. One way analysis of variance (in some cases for repeated measures) was followed by Bonferroni post-hoc analysis. 2-way analysis of variance (ANOVA) for repeated measures (rmANOVA) was followed by Bonferroni post-hoc analysis. Non-parametric analysis was carried out using the Mann-Whitney *U* test. Contingency tables were analyzed using *χ*^2^ test if the tables were of sufficient size, otherwise the Fisher’s exact test was used. A *p*<0.05 was considered statistically significant. First, group differences were verified by ANOVA, followed - if appropriate - by post-hoc tests which considered differences between HAB vs. NAB or LAB vs. NAB. As the manifestation of *high-anxiety* and *low-anxiety* phenotypes via selective breeding most likely involved different complex multigenic changes, a direct comparison of HAB against LAB is inappropriate. Therefore we only compared HAB and LAB to the common NAB control.

## Author Contributions

AJG conceptualization, data curation, formal analysis, investigation, methodology, project administration, software, supervision, visualization, writing - original draft preparation; NA investigation, visualization; SAB investigation; PK formal analysis; DEH investigation; MN resources; ME investigation; SFK methodology, investigation; CJR methodology, investigation; BTB methodology, investigation, visualization; MC resources, software, supervision, visualization, writing - review & editing; CTW formal analysis, methodology, project administration, resources, supervision, validation, writing - original draft preparation, review & editing.

## Supporting information

### Supplemental Movie 1

Movie explaining the Robocat task also known as the *Panic Box*.

**Supplemental Figure 1.**
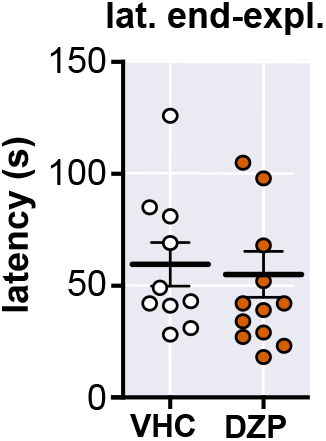
Unaltered latency for end-exploration in the IndyMaze task with DZP treatment.

**Supplemental Figure 2.**
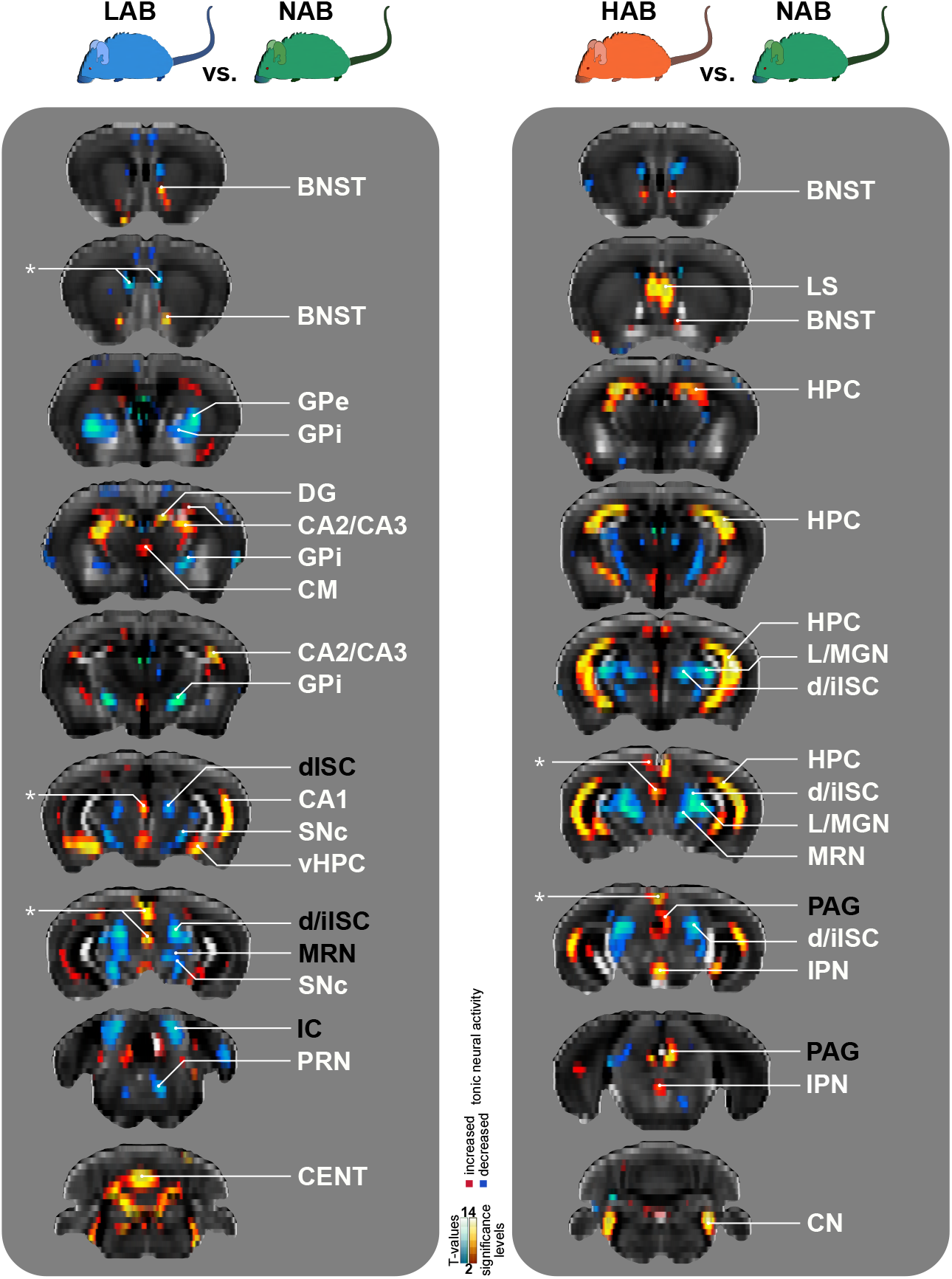
Complete MEMRI Data Set. Asterisks indicate potential artifacts which have been observed to occur close to the brain surface or the ventricular system. **BNST** bed nucleus of the stria terminalis, **CA1-3** cornu ammonis 1-3, **CENT** central lobule of the cerebellum, **CM** central medial nucleus of the thalamus, **CN** cochlear nucleus, **DG** dentate gyrus, **d/ilSC** deep/intermediate layers of the superior colliculus, **GPe** globus pallidus external segment, **GPi** globus pallidus internal segment, **HPC** hippocampus proper, **IPN** interpedunculopontine nucleus, **LS** lateral septal nucelus, **L/MGM** lateral/medial geniculate, **MRN** midbrain reticular nucleus, **PAG** periaqueductal gray, **PRN** pontine reticular nucleus, **SNc** substantia nigra pars compacta, **vHPC** ventral portion of the hippocampus proper.

**Supplemental Figure 3.**
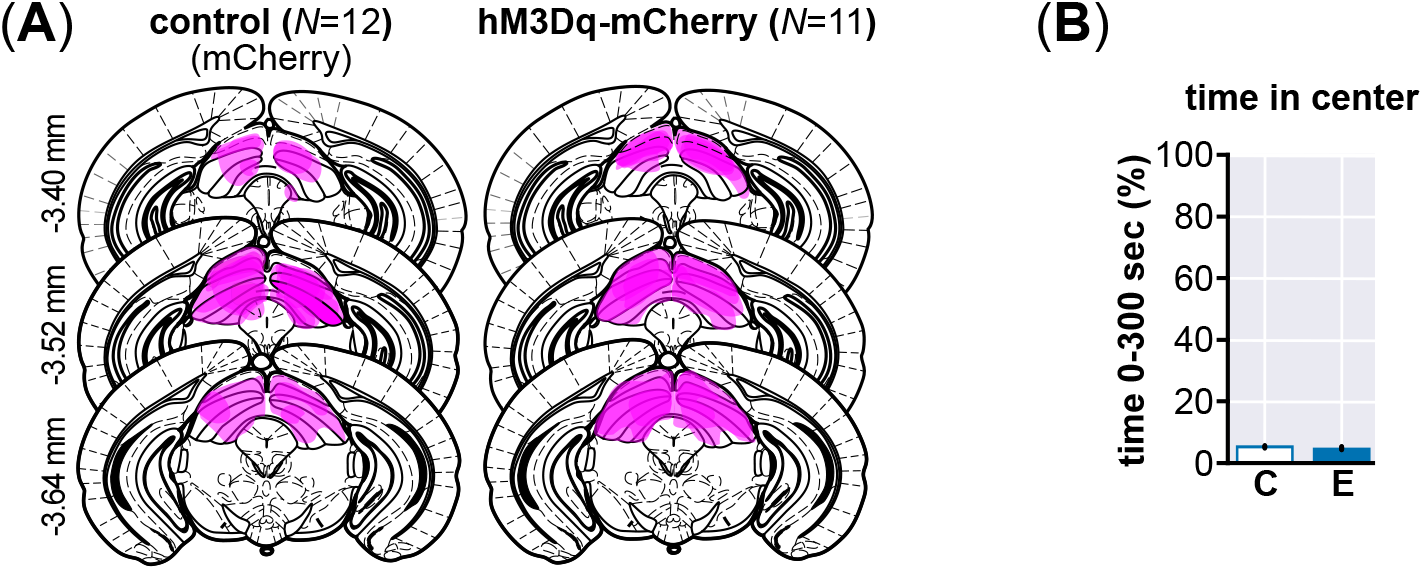
Summary of histological analysis of virus spread in LAB animals and aditional EPM measures.

**Supplemental Figure 4.**
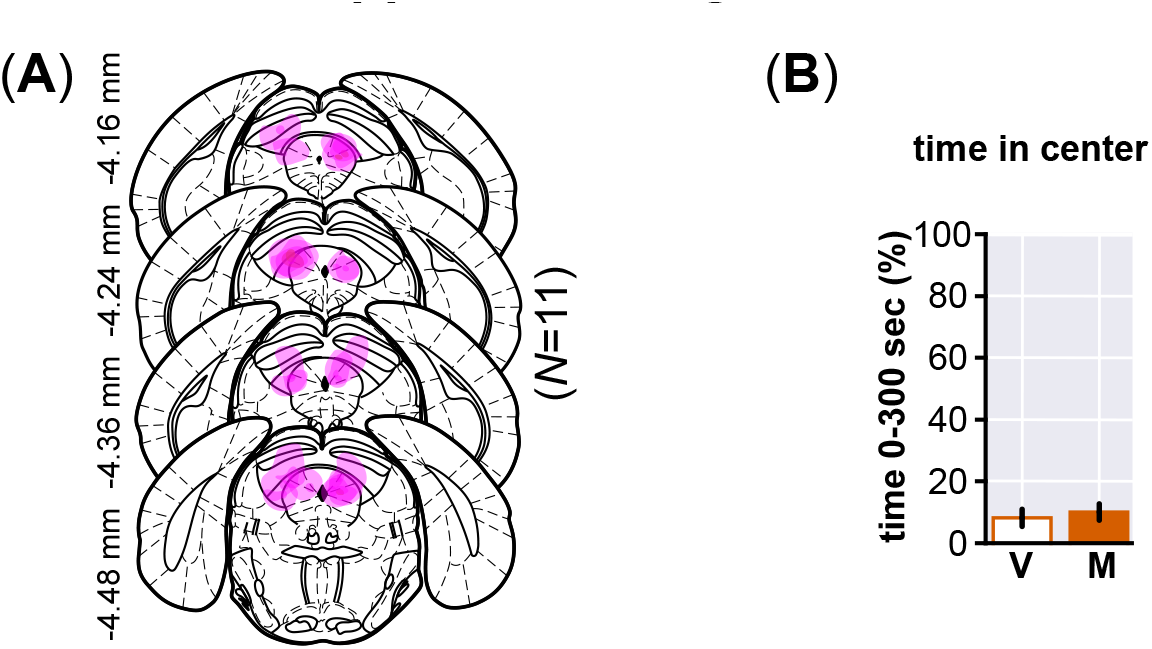
Summary of histological analysis of MUSC spread in HAB animals and aditional EPM measures.

